# Histone H2B Monoubiquitylation Regulates Elongation-to-Termination Transition in RNA Polymerase II Transcription

**DOI:** 10.1101/2024.07.27.605434

**Authors:** Tirthankar Bandyopadhyay, Baidehi Basu, Pabitra K. Parua

## Abstract

RNA Polymerase II (Pol II)-dependent transcription is meticulously regulated during initiation, elongation, termination, and two pivotal transitions: from initiation to elongation and from elongation to termination ^1^. The successful transition from elongation to termination is vital for accurate transcription termination ^2,3^. While elongation and termination have been extensively studied, the regulatory mechanisms governing the transition between these stages are not fully understood. Here we report that, in fission yeast *Schizosaccharomyces pombe* (*S. pombe*) and human cells, Cdk9 and its cyclin partner (Cyclin T in humans, and Pch1 in *S. pombe*) disengage from the elongation machinery near gene 3’-ends, as Pol II crosses the cleavage and polyadenylation signal (CPS). Concurrently, chromatin immunoprecipitation (ChIP) data shows an increased association of Dis2 (the PP1 ortholog in *S. pombe*)^4^ beyond the CPS. Additionally, ChIP-seq indicates that histone H2B monoubiquitylation (H2Bub1) is pivotal for Cdk9 occupancy but serves differential roles along the gene. H2Bub1 facilitates Cdk9 chromatin recruitment during elongation but prompts Cdk9 dissociation from the elongation complex as Pol II traverses the CPS. Furthermore, in *S. pombe*, disruptions of H2Bub1 in *htb1-K119R* ^5^ and sustained levels in *ubp8D* ^6^ inversely affect the chromatin occupancy of Pol II, Pol II CTD phosphorylation within the heptad repeats, Spt5, phosphorylated Spt5 (pSpt5), Dis2, components of the mRNA 3’-end processing complex (Pfs2 and Pla1), and termination factors Rhn1 (Rtt103) and Pcf1 around the CPS— diminishing with H2Bub1 loss and increasing with H2Bub1 stabilization. In essence, our data indicates that the H2Bub1-mediated eviction of Cdk9 as Pol II navigates the CPS precipitates a rise in PP1 binding, which, in turn, dephosphorylates pSpt5, slowing Pol II to facilitate efficient termination. The unique modulation of Cdk9 occupancy by H2Bub1 during transcription progression presents a compelling, novel concept requiring further exploration.

---

The elongation to termination transition in RNA Polymerase II (Pol II)-dependent transcription is regulated by a switch mechanism involving cyclin-dependent kinase 9 (Cdk9) and protein phosphatase 1 (PP1) that modulates the phosphorylation of Spt5 (Fig. 1a) ^7^. Yet, the precise mechanisms overseeing the activity switch of Cdk9 and PP1, as Pol II transits through the cleavage and polyadenylation sequence (CPS), remain undefined. While investigating the mechanism of timely reversal of Cdk9-PP1 activity (Fig. 1a) we hypothesized that either Cdk9 becomes inactive or dissociates from the elongation complex as Pol II transits through the CPS. To understand the distribution of Cdk9 and its cyclin partner cyclin T1 (CycT1; Pch1, ortholog of CycT1 in fission yeast) we conducted ChIP experiments in *Schizosaccharomyces pombe* (*S*. *pombe*). The ChIP-qPCR results showed a gradual decrease of the chromatin occupancy of Cdk9 and Pch1 towards the 3’-end of *psu1^+^*, *pgk1^+^*, and *pyk1^+^*—more predominant corresponding to the amplicon positioned beyond the CPS of these genes (Extended Data Fig. 1a-c). The ChIP-seq browser tracks indicate a diminished level of Cdk9 and Pch1 beyond the CPS of *exg1^+,^ eno101^+^*, *pma1^+^*and *act1^+^* (Fig. 1b, Extended Data Fig. 1d). Wherever the chromatin occupancy of Cdk9 and Pch1 dropped, Pol II and Spt5 accumulated, but pSpt5 reduced (Fig. 1b). The metagene plots further supported the notion of a global decrease in the chromatin occupancy of Cdk9 and Pch1 as Pol II approaches the 3’-end of genes, with a pronounced reduction beyond the CPS (Fig. 1c,d; Extended Data Fig. 1e,f). Similar to protein-coding genes, the occupancy of Cdk9 and Pch1 decreases at the 3’-end of small nucleolar RNA (snoRNA) coding genes (Extended Data Fig. 1g).

**Figure 1.**
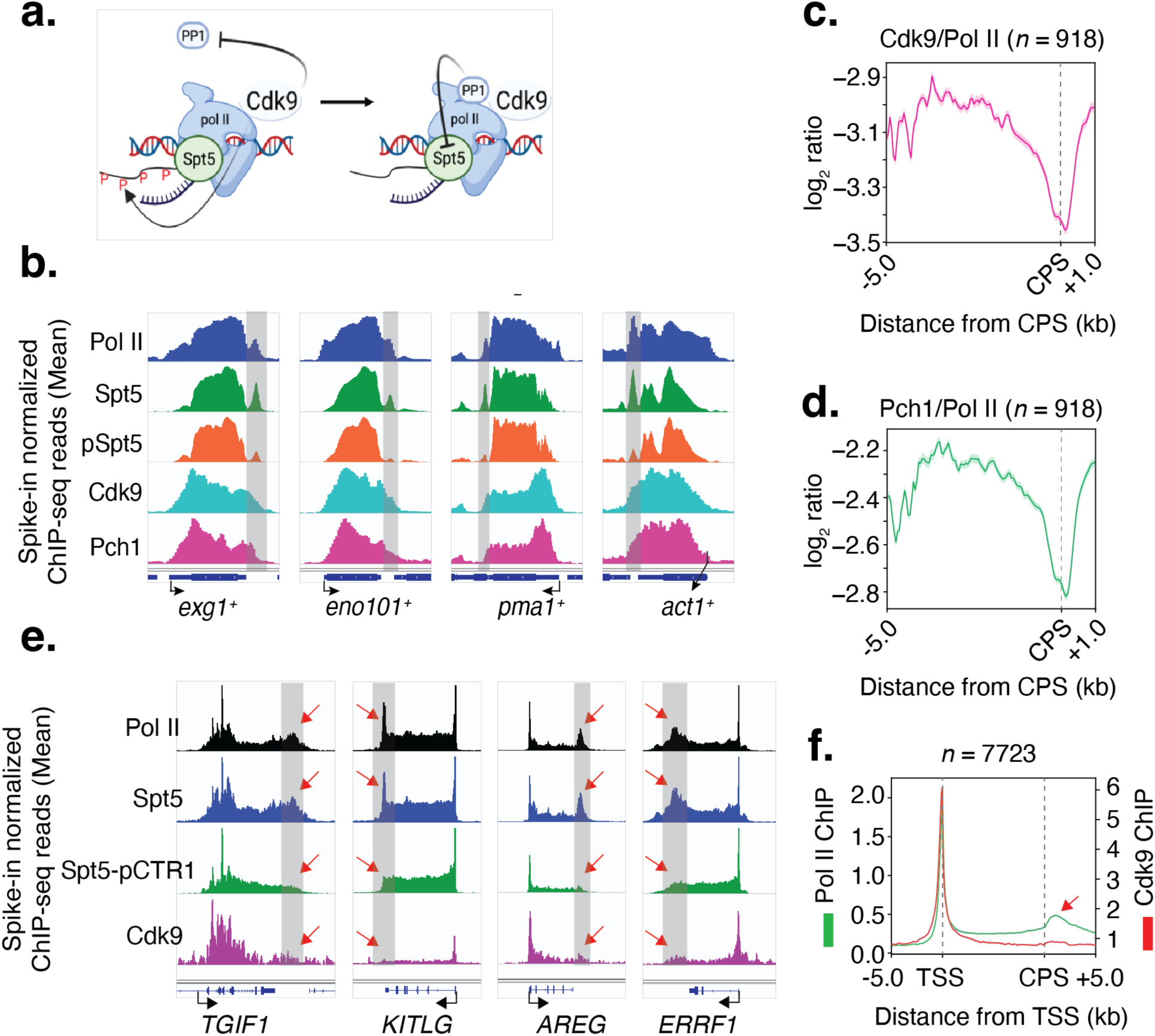
Cdk9 and Pch1 dissociate from the elongation complex when Pol II passes the CPS. **a**, The model shows the Cdk9-PP1 switch controlling the transition from elongation to termination (PP1, protein phosphatase 1). The figure is prepared using (https://www.biorender.com/). **b**, In fission yeast, Cdk9 and Pch1 leave the elongation complex during the transition to termination. Individual gene tracks show that Cdk9 and Pch1 occupancy drops downstream of the CPS at the Pol II 3’-pause site (indicated by a gray shade). At the same location, Spt5 accumulates while Spt5 phosphorylation (pSpt5) decreases. **c**, **d**, Genome-wide distribution of Cdk9 and Pch1 on 1 kb separated genes (*n* = 918). Metagene analyses reveal a sharp drop in Cdk9 and Pch1 occupancy when Pol II approaches the CPS (marked by a vertical dotted line). Genes were sorted based on region length between -5 kb to +1 kb relative to the CPS. The mean log_2_ ratio over Pol II was plotted in the summary image. **e**, In humans, Cdk9 releases from the elongation complex after the CPS. Representative gene tracks show that Cdk9 occupancy decreases downstream of the CPS where the Pol II 3’-pause peak is positioned (indicated by a gray shade). At the same location, Spt5 accumulates while Spt5-CTR1 phosphorylation (Spt5-pCTR1) decreases. Pol II, Spt5, and Spt5-pCTR1 ChIP data utilized in this analysis were obtained from a published study ^42^. Peaks are indicated by a red arrow. **f**, Genome-wide distribution of Cdk9 and Pol II on 10 kb separated genes (*n* = 7723). Metagene analyses reveal reduced levels of Cdk9 occupancy downstream of the CPS where Pol II accumulates (indicated by a red arrow). Genes were sorted based on Pol II occupancy. (Note: In this representation, the regions between the TSS and the CPS have been scaled to enable comparisons among genes of different lengths.) The TSS and CPS are marked by vertical dotted lines.

Similar to *S. pombe*, in human cells, the ChIP-qPCR distributions of Cdk9 across *cMYC, ERRFI1*, *GAPDH* and *VEGFA* showed the gradual reduction of Cdk9 occupancy towards the 3’- end of those genes (Extended Data Fig. 2a-d). The ChIP-seq browser tracks of four representative genes (*TGIF1*, *KITLG*, *AREG*, and *ERRFI1*) indicated diminished levels of Cdk9 beyond the CPS, where Pol II and Spt5 peaked but the Spt5 CTR1 (C-terminal repeat region 1) phosphorylation (Spt5-pCTR1) dropped (Fig. 1e, Extended Data Fig. 2e). Furthermore, metagene plots demonstrated a global decrease in Cdk9 chromatin occupancy beyond the CPS at location corresponding to the 3’-paused Pol II (Fig. 1f, Extended Data Fig. 2f-h). Overall, these findings suggested that Cdk9 dissociated from the elongation complex as Pol II traverses the CPS and this phenomenon is conserved in *S. pombe* and human.

Previous studies have shown that during elongation, Cdk9-mediated phosphorylation inhibits the activity of Dis2, a PP1 ortholog in *S. pombe*, and reduces its chromatin occupancy ^7^. In this study, we demonstrate the dissociation of Cdk9 from the elongation complex as Pol II traverses the CPS. Drawing from consistent findings in both *S. pombe* and human studies, we propose that the disengagement of Cdk9 and its partner CycT (Pch1) from the elongation complex leads to increased chromatin occupancy of Dis2. The ChIP-qPCR analysis in *S. pombe* indicates a dramatic incline in the occupancy of Dis2 beyond the CPS of the representative genes, *psu1^+^*, *pgk1^+^* and *pyk1^+^* (Fig. 2a-c, Extended Data Fig. 3a-c). The ChIP-seq browser tracks for the sample genes display a significant buildup of Dis2 past the CPS, aligning with the location of the 3’-paused Pol II (Fig. 2d, Extended Data Fig. 3d). Metagene analyses further indicated a gradual increase in Dis2 occupancy towards the 3’-end of Pol II-transcribed genes, with a sharp peak beyond the CPS (Fig. 2e, Extended Data Fig. 3e). To the best of our knowledge, this is the first report detailing the genome-wide distribution of Dis2 (PP1), demonstrating significant accumulation of Dis2 beyond the CPS. Overall, our results suggested an inverse relationship between Cdk9 and Dis2: as the former leaves the elongation complex, the latter begins to accumulate. This accumulation likely explains the Dis2-mediated rapid dephosphorylation of pSpt5 beyond the CPS ^7^, which in turn appears to slow Pol II, promoting faithful termination. However, the question remains: What triggers the eviction of Cdk9 from the elongation complex during the transition from elongation to termination?

**Figure 2.**
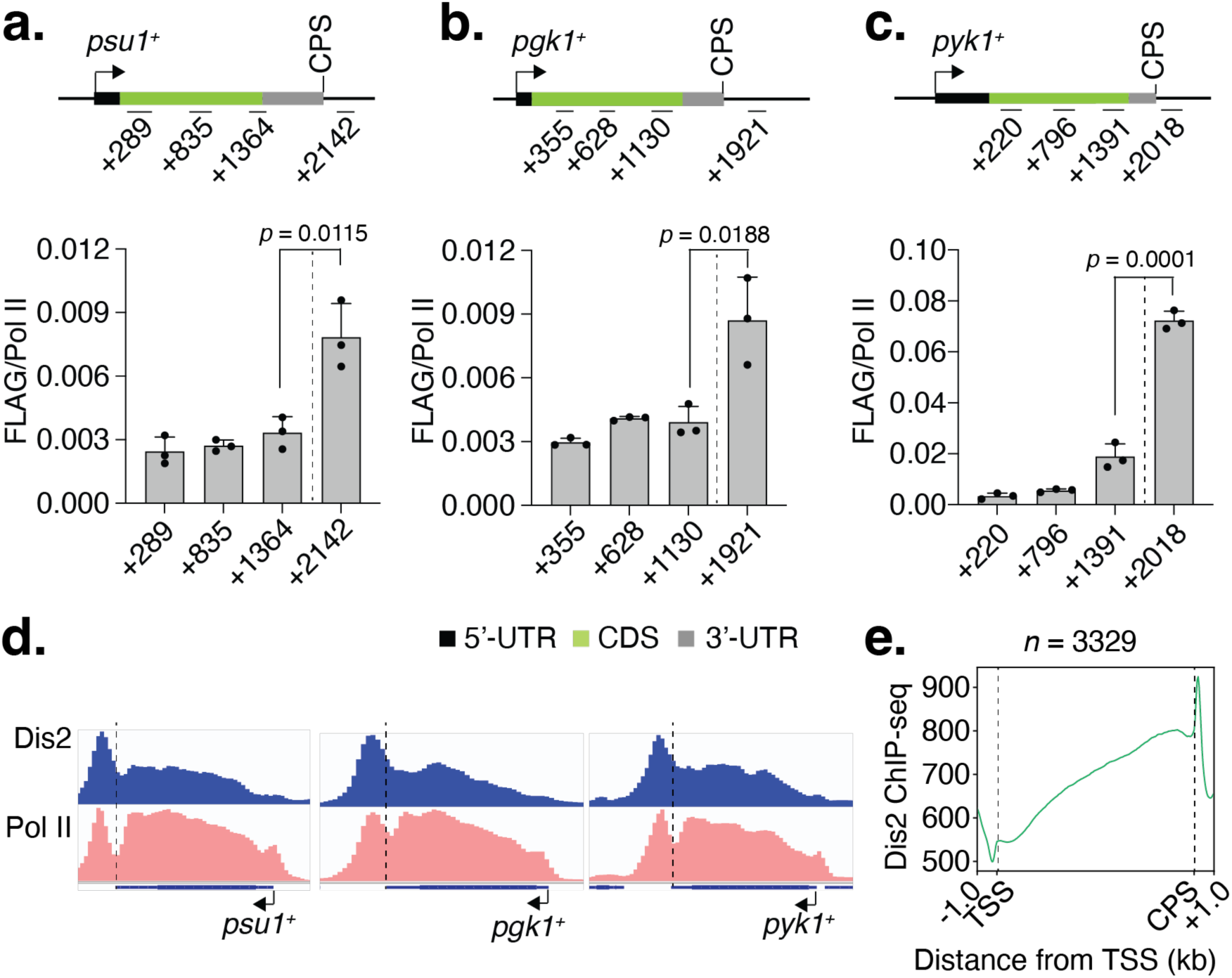
Dis2 accumulates beyond the CPS. **a-c**, Dis2 ChIP was performed using chromatin extract from *S. pombe* cells expressing Dis2-FLAG, which is a C-terminal FLAG-tagged version of Dis2. ChIP-qPCR analyses show the distribution of Dis2 on *psu1^+^*, *pgk1^+^*, and *pyk1^+^*genes. The results reveal a sharp increase in Dis2 occupancy beyond the CPS at the location of the fourth amplicon. In the schematic diagram of genes, the 5’-untranslated region (5’-UTR), the coding sequence (CDS), and the 3’-UTR are demarcated by black, green, and gray, respectively. The regions of the genes scanned are indicated by amplicon positions. The experiments were performed with three biological replicates; one-sided error bars show standard deviation (s.d.) of mean; *p*-values (Student’s *t-*test) are indicated in the plots between third (upstream of the CPS) and fourth (downstream of the CPS) amplicons. **d**, ChIP-seq browser tracks show the distribution of Dis2 and Pol II across *psu1^+^*, *pgk1^+^*, and *pyk1^+^* genes. The results show accumulation of Dis2 after the CPS corresponding to the position where Pol II accumulates. **e**, Metagene analyses show a sharp accumulation of Dis2 beyond the CPS of Pol II-transcribed genes (*n* = 3329). In this plot, the regions between the TSS and the CPS have been scaled to enable comparisons among genes of different lengths. Vertical dotted lines mark the TSS and CPS.

Earlier studies identified the functional interconnection between Cdk9 and the mono- ubiquitylation of histone H2B (H2Bub1) at K120 in humans (K123 in *S. cerevisiae*, and K119 in *S. pombe*) ^8^. A decrease in Cdk9 activity negatively impacts the stability of H2Bub1, partly through the phosphorylation of Spt5, which facilitates the formation of H2Bub1 ^9–14^. In a feedback scenario, H2Bub1, which is highly enriched in the gene bodies of active genes and sharply decreases after the CPS ^15,16^, also controls Cdk9 recruitment in *S. pombe* ^8^ and human cells ^17,18^. The loss of H2Bub1 levels decreases chromatin occupancy of Cdk9 on gene body, although this has been explored only at a limited number of loci. We propose that H2Bub1 preserves the chromatin occupancy of Cdk9 during the elongation phase of transcription, and the loss of H2Bub1 as Pol II traverses the CPS might trigger the disengagement of Cdk9 from the elongating Pol II complex. The ChIP-seq browser tracks showed that across the genes *act1^+^*, *exg1^+^*, and *eno101^+^*, disruption of H2Bub1 led to a decrease in Cdk9 occupancy at the 5’- end and within the gene body (Fig. 3a, Extended Data Fig. 4a). However, upstream and downstream of the CPS, the Cdk9 occupancy was increased in *htb1-K119R* cells compared to wildtype. The metagene plot showed a sharp increase in Cdk9 occupancy around the CPS upon loss of H2Bub1 (Fig. 3b, Extended Data Fig. 4b). The upstream increase of Cdk9 occupancy indicates H2Bub1 rearranges Cdk9 association with the elongation complex before Pol II passes through the CPS. These results suggest that H2Bub1 differentially impacts the chromatin occupancy of Cdk9 across a transcription cycle. The decrease in Cdk9 occupancy upon loss of H2Bub1 at the 5’-end to gene body is consistent with the previous findings determined by ChIP-qPCR for a few genes ^8^. However, the effects of H2Bub1 on Cdk9 occupancy at the 3’-end of genes have not been examined before. These potential paradigm-shifting roles of H2Bub1 in regulating Cdk9’s occupancy differentially across a gene is intriguing and warrant further investigation.

**Figure 3.**
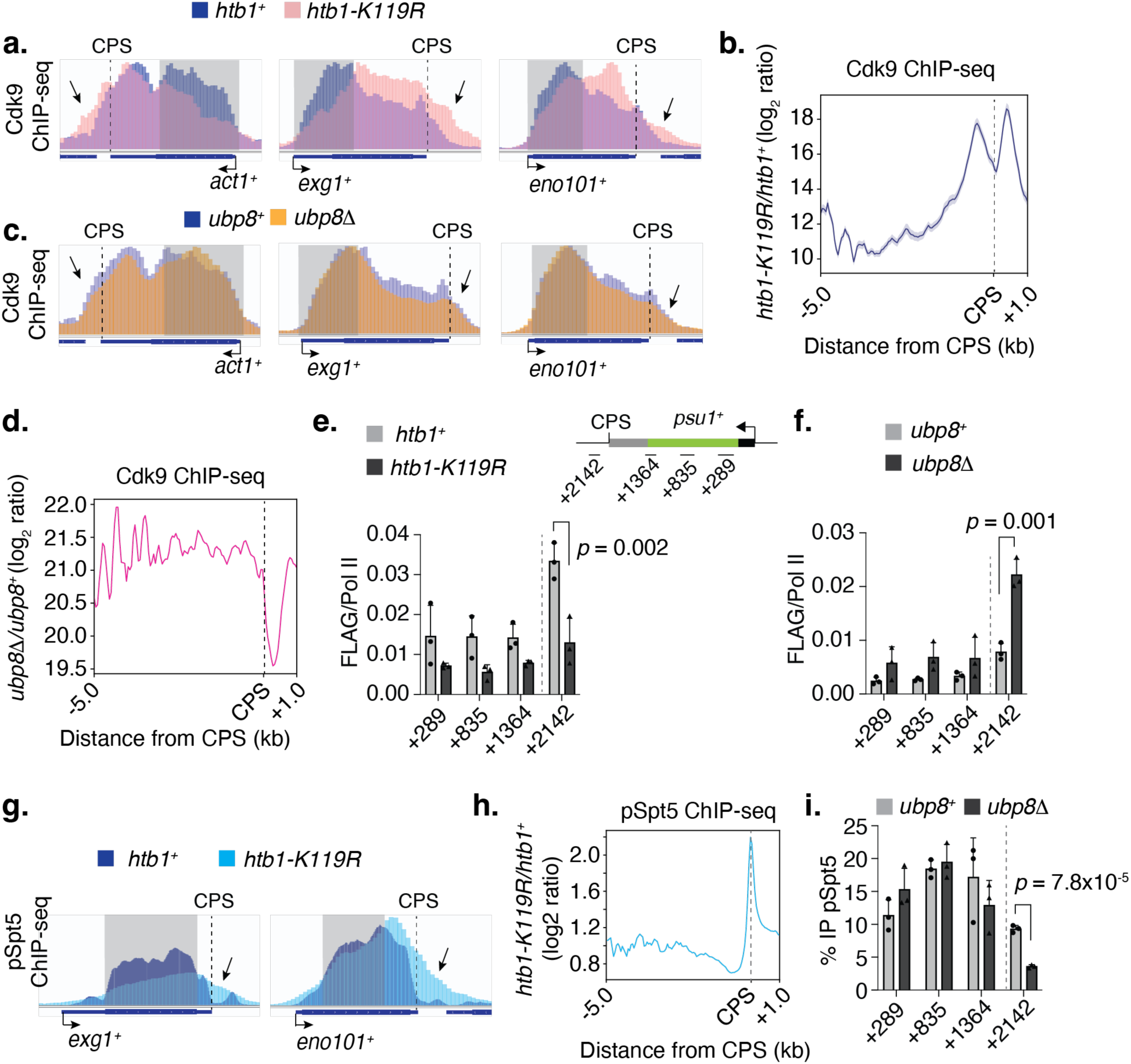
H2Bub1 regulates Cdk9’s chromatin occupancy. **a**, H2Bub1 differentially regulates Cdk9’s chromatin occupancy throughout the transcription cycle. Individual browser tracks display an increase in Cdk9 occupancy downstream of the CPS (marked by an arrow) and a decrease at the 5’ end (marked by a gray shaded area) of the example genes upon loss of H2Bub1. Cdk9 ChIP-seq was performed in cells expressing 13Myc-tagged Cdk9 in the *htb1^+^* and *htb1-K119R* backgrounds using an anti-Myc antibody. Blue and pink tracks represent the Cdk9 distribution in *htb1^+^* and *htb1-K119R* cells, respectively, overlaid. **b**, The metagene plot shows the genome-wide distribution of Cdk9 on 1 kb separated genes (*n* = 918) from -5.0 to +1.0 kb relative to the CPS. Results indicate accumulation of Cdk9 upstream and downstream of the CPS in *htb1-K119R* cells. The log_2_ ratio of mean ChIP-seq values in *htb1-K119R* over *htb1^+^* is plotted. **c**, Similar to panel **a**, but Cdk9-13Myc ChIP was conducted in *ubp8^+^* and *ubp8Δ* cells. Browser tracks reveal minimal changes in Cdk9 distribution at the 5’ end (indicated by a gray shaded area) of genes, but a decrease beyond the CPS (indicated by an arrow) in *ubp8Δ* cells. Blue and orange tracks represent the Cdk9 distribution in *ubp8^+^*and *ubp8Δ* cells, respectively, overlaid. **d**, The metagene plot shows the genome-wide distribution of Cdk9 on 1 kb separated genes (*n* = 918) from -5.0 to +1.0 kb relative to the CPS. Results indicate a rapid drop of Cdk9 downstream of the CPS. The log_2_ ratio of mean ChIP-seq values in *ubp8Δ* over *ubp8^+^*is plotted. **e**, ChIP-qPCR analyses show the distribution of Dis2-FLAG on the *psu1^+^* gene in *htb1^+^* (gray bar) and *htb1-K119R* (black bar) cells. Results indicate a significant decrease in Dis2 occupancy after the CPS in *htb1-K119R* cells. **f**, ChIP-qPCR analyses show the distribution of Dis2-FLAG on the *psu1^+^* gene in *ubp8^+^* (gray bar) and *ubp8Δ* (black bar) cells. Results indicate a significant increase in Dis2 occupancy after the CPS in *ubp8Δ* cells. For **e**, **f**, experiments were performed in cells expressing Dis2 as C-terminal FLAG tagged; graphs plotted the ratio of Dis2-FLAG ChIP over Pol II along the Y-axis. **g**, Similar to panel **a**, but pSpt5 ChIP-seq was conducted in *htb1^+^*and *htb1-K119R* cells. Individual browser tracks display an increase in pSpt5 occupancy downstream of the CPS (marked by an arrow) and a decrease at the 5’ end (marked by a gray shaded area) of the example genes upon loss of H2Bub1. Blue and cyan tracks represent the pSpt5 distribution in *htb1^+^* and *htb1-K119R* cells, respectively, overlaid. **h**, The metagene plot shows the genome-wide distribution of pSpt5 on 1 kb separated genes (*n* = 918) from -5.0 to +1.0 kb relative to the CPS. Results indicate accumulation of pSpt5 around the CPS in *htb1-K119R* cells. The log_2_ ratio of mean ChIP-seq values in *htb1-K119R* over *htb1^+^* is plotted. **i**, ChIP-qPCR analyses show the distribution of pSpt5 on the *psu1^+^* gene in *ubp8^+^* (gray bar) and *ubp8Δ* (black bar) cells. Results indicate a significant decrease in pSpt5 occupancy after the CPS in *ubp8Δ* cells. For panels **e**, **f**, and **i**, experiments were performed with three independent biological replicates (*n* = 3); one-sided error bars show the standard deviation (s.d.) of mean ChIP values; *p*-values (Student’s t-test) are indicated in the plots. For panels **a**-**i**, vertical dotted lines mark the CPS.

Should H2Bub1 regulate Cdk9 occupancy during the elongation to termination transition, we anticipated observing reversal of H2BK119R-phenotype in distribution of Cdk9 when H2B undergoes hyper-monoubiquitylation. To examine this, we checked Cdk9 distribution in *ubp8D* cells by ChIP-seq. Ubp8 serves as a ubiquitin C-terminal hydrolase in the SAGA complex and its absence fortifies H2Bub1 at K119. In *ubp8D* cells, the ChIP-seq browser tracks showed a reduced level of Cdk9 towards the 3’-end of the genes *act1^+^*, *exg1^+^*, and *eno101^+^*(Fig. 3c, Extended Data Fig. 4a). The metagene plot revealed no accumulation of Cdk9 upstream of the CPS; however, a sharp and widespread decline in Cdk9 occupancy beyond the CPS upon Ubp8 loss (Fig. 3d, Extended Data Fig. 4c). These observations reinforced our hypothesis that hyper- monoubiquitylation of H2B, situated upstream of the CPS ^16^, aids in dislodging Cdk9 from the elongation complex as it traverses the CPS. Accordingly, the loss of H2Bub1 engages Cdk9 around the CPS.

We identified an inverse relationship between Cdk9 and Dis2 distribution throughout the transcription cycle. We expanded our studies to determine whether H2Bub1 inversely regulates Dis2 occupancy relative to Cdk9. To investigate this, we examined Dis2 occupancy in *htb1- K119R* and *ubp8Δ* cells using ChIP. The ChIP-qPCR results indicate that the loss of H2Bub1 decreases Dis2 occupancy, specifically beyond the CPS of three genes, *psu1+*, *pgk1^+^* and *pyk1^+^* (Fig. 3e; Extended Data Fig. 4d,e, top panel). Conversely, persistent levels of H2Bub1 increase Dis2 occupancy in the same region (Fig. 3f; Extended Data Fig. 4d,e, bottom panel). In *htb1- K119R* cells, consistent with the alterations in Cdk9 and Dis2 occupancy, ChIP-seq browser tracks show an increase in Spt5 phosphorylation (pSpt5) around the CPS and a decrease at the 5’-end of genes, respectively (Fig. 3g, Extended Data Fig. 5a,b). However, in agreement with previous reports, Spt5 phosphorylation decreases in whole-cell extracts following the loss of H2Bub1 (in *htb1-K119R* cells), whereas it increases upon stabilization of H2Bub1 in *ubp8Δ* cells ^8^ (Extended Data Fig. 5c). Presumably, the loss of Spt5 phosphorylation from the 5’-end to the middle of the gene body seen in *htb1-K119R* cells accounts for the global decrease in whole-cell extracts. Metagene analyses reveal a sharp accumulation of pSpt5 levels around the CPS in *htb1-K119R* cells (Fig. 3h, Extended Data Fig. 5d). However, in *ubp8Δ* cells, pSpt5 decreases around the CPS (Fig. 3i, Extended Data Fig. 5e,f), agreeing with the accumulation of H2Bub1 upstream of the CPS, where pSpt5 decays progressively ^16^. Overall, our results demonstrate that H2Bub1 inversely regulates Cdk9 and Dis2 occupancy during the transition from elongation to termination, leading to the modulation of Spt5 phosphorylation.

The transition from elongation to termination is closely linked with pre-mRNA 3’-end cleavage and polyadenylation. Once Pol II transcribes the cleavage and polyadenylation sequence, the cleavage and polyadenylation factor (CPF), a multiprotein complex, is recruited. This in turn facilitates the 3’-end cleavage and polyadenylation of pre-mRNA ^19^. We sought to determine whether the cleavage and polyadenylation of pre-mRNA triggers the dissociation of Cdk9 from the elongation complex. To investigate this, we conducted Cdk9 ChIP in *pfs2-11* cells. Since Pfs2 is a crucial component of the CPF complex, the *pfs2-11* mutant shows a deficiency in mRNA 3’-end processing at 37°C ^20^. ChIP-qPCR analysis demonstrated that there were no changes in Cdk9 distribution on representative genes following the heat inactivation of Pfs2 (Extended Data Fig. 6a,b). This finding aligns with previous reports showing no changes in CPF occupancy when Cdk9 is inhibited and no alterations in chromatin-bound levels of Spt5 phosphorylation upon Pfs2 heat inactivation ^7^. However, under these same conditions, Pol II occupancy increased at the 3’-end of the genes, suggesting termination defects. These findings suggest that: 1) the productive cleavage and polyadenylation event is not the trigger for Cdk9’s dissociation from the elongation complex; it is either independent of or occurs prior to mRNA cleavage and polyadenylation, and 2) the accumulation of Cdk9 upon loss of H2Bub1 is not due to the accumulation of Pol II.

These results invoked us to determine whether H2Bub1 regulates the recruitment of CPF complex. To investigate that we conducted ChIP-qPCR for Pla1, mRNA cleavage and polyadenylation specificity factor poly(A) polymerase, and Pfs2 in *htb1-K119R* and *ubp8Δ* cells. The ChIP-qPCR results show a decrease or an increase or in both Pfs2 (Extended Data Fig. 6c,d) and Pla1 (Extended Data Fig. 6e,f) recruitment beyond the CPS of the representative genes when H2Bub1 is disrupted (in *htb1-K119R* cells) or persistent (in *ubp8Δ* cells), respectively. These results indicate that H2Bub1 facilitates the recruitment of CPF complex. Moreover, these effects of H2Bub1 on the occupancy of these two factors of the 3’-end processing complex, are opposite of the effects on the occupancy of Cdk9 at the similar regions. Additionally, our results imply that H2Bub1-mediated Cdk9 eviction happens upstream of 3’-end processing.

Previous studies report that PP1-mediated Spt5 dephosphorylation facilitates Pol II slowing beyond the CPS ^7^. This slow Pol II favors faithful termination by allowing the recruitment of termination factors. In our current study, we found that the loss of H2Bub1: 1) leads to accumulation of Cdk9 around the CPS, 2) disrupts the chromatin occupancy of Dis2, subsequently, 2) increases the Spt5 phosphorylation beyond the CPS, and 3) decreases the association of cleavage and polyadenylation factors with the elongation complex, which is required for proper 3’-end processing of mRNA. Given that Spt5 dephosphorylation and proper 3’-end processing are essential for faithful termination, we aimed to investigate whether H2Bub1 regulates termination. To examine this, we conducted Pol II ChIP-seq in *htb1-K119R* and *ubp8Δ* cells. In *htb1-K119R* cells, the ChIP-seq bowser tracks showed a wider distribution of Pol II past the CPS, indicating readthrough transcription (Fig. 4a,b, Extended Data Fig 7). Conversely, in cells lacking Ubp8 (*ubp8Δ*), there was a tighter distribution of Pol II around the termination region, which suggests a more effective termination process (Fig. 4a,b, Extended Data Fig 7).

**Figure 4.**
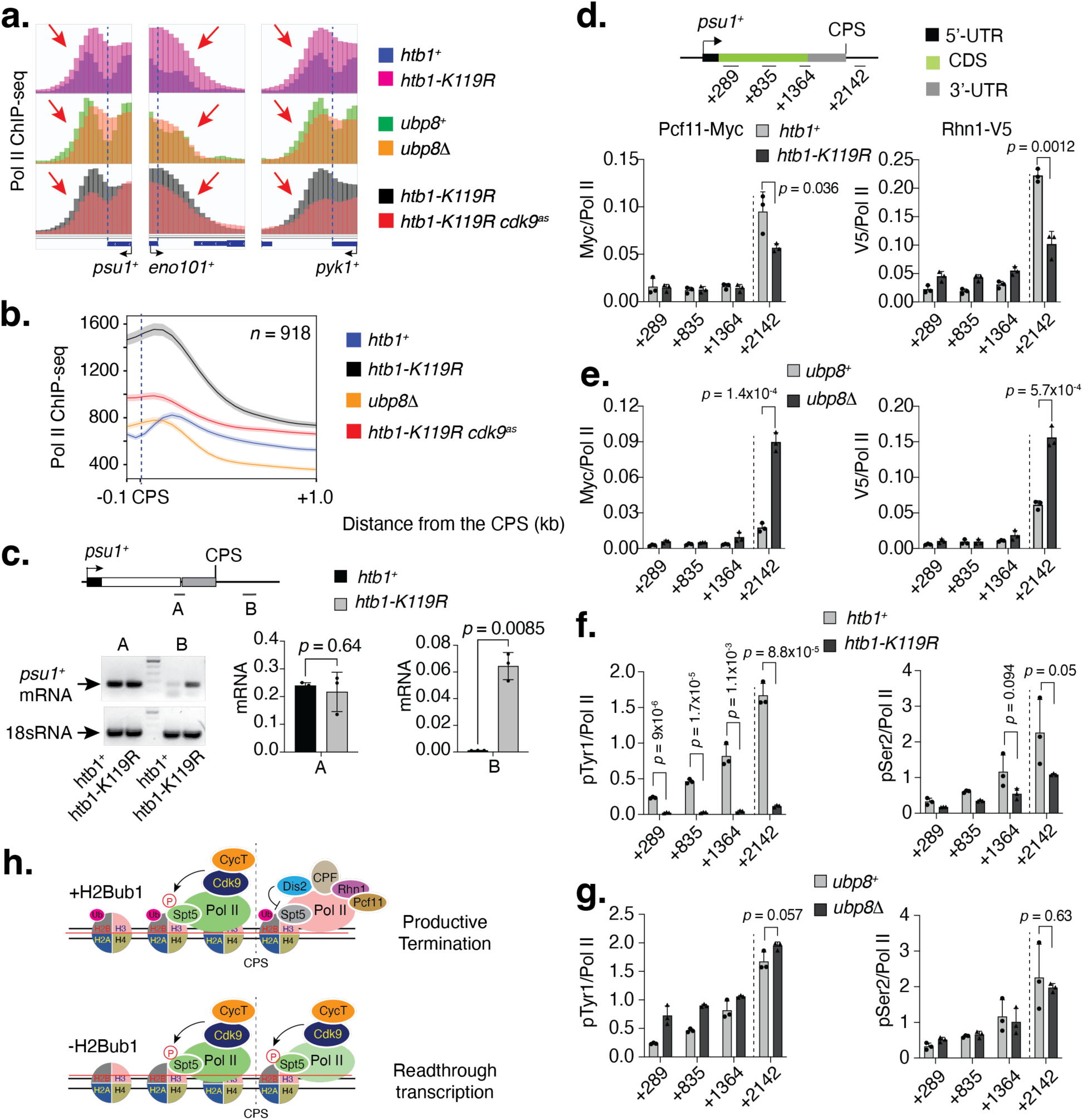
H2Bub1 controls termination. **a**, Loss of H2Bub1 impairs termination. ChIP-seq browser tracks reveal an extended Pol II distribution in *htb1-K119R* cells beyond the regular termination zone seen in *htb1^+^* cells, indicating readthrough transcription, i.e., termination defects. The blue and pink tracks, representing the Pol II distribution in *htb1^+^* and *htb1-K119R* cells, respectively, are overlaid. Conversely, persistent levels of H2Bub1 favor termination. ChIP-seq browser tracks show a narrowed Pol II distribution in the termination zone in *ubp8Δ* cells, indicating that hyper mono-ubiquitylation favors termination. The green and orange tracks, representing the Pol II distribution in *ubp8^+^* and *ubp8Δ* cells, respectively, are overlaid. Cdk9 inhibition in the *htb1-K119R* background narrows Pol II distribution, suggesting a rescue of termination defects. This indicates that the accumulation of Cdk9 following the loss of H2Bub1 is the main trigger for termination defects by continuing elongation. The black and red tracks, representing the Pol II distribution in *htb1-K119R*, with or without inhibition of Cdk9, respectively, are overlaid. Pol II ChIP-seq data following Cdk9 inhibition in *htb1-K119R* cells, used in this analysis, were obtained from a published study ^16^. The regions of alterations in Pol II distribution after the CPS are indicated by red arrows. The vertical dotted lines mark the CPS. **b**, The metagene plot shows the genome-wide distribution of Pol II on 1 kb separated genes (*n* = 918) from -0.1 kb to +1.0 kb relative to the CPS. Results indicate a broadening of Pol II distribution in the termination zone in *htb1-K119R* cells, implying termination defects. Conversely, in *ubp8Δ* cells, the Pol II distribution narrowed compared to wildtype (*ubp8^+^*) cells, indicating premature termination. Cdk9 ^16^ inhibition in the *htb1-K119R* background narrows Pol II distribution, indicating a rescue of termination defects. The log_2_ ratio of mean ChIP-seq values in *htb1- K119R* (or *ubp8Δ*) with active or inactive Cdk9 over wildtype (*htb1^+^*/*ubp8^+^*) is plotted. **c**, The left panel shows an agarose gel visualizing transcript production from the upstream (A) and downstream (B) regions of the CPS of the *psu1^+^* gene. The results indicate the synthesis of *psu1^+^*mRNA beyond the regular termination zone in *htb1-K119R* cells, with little to no mRNA synthesis in *htb1^+^* cells. The right panel shows RT-qPCR analyses quantifying *psu1^+^* mRNA production from the same regions, A and B, indicating increased mRNA synthesis beyond the termination zone in *htb1-K119R* cells. For panel **c**, individual data points shown in the plots were obtained from three independent biological replicates (*n* = 3); two-sided error bars represent the standard deviation (s.d.) of mean qRT-PCR values; *p*-values (Student’s t-test) are indicated in the plots. **d**, Loss of H2Bub1 impairs the recruitment of termination factors. ChIP-qPCR analyses reveal a significant decrease in the recruitment of termination factors, Pcf11-Myc and Rhn1-V5, after the CPS of *psu1^+^*in *htb1-K119R* cells. **e**, Persistent levels of H2Bub1 facilitate termination factor recruitment. ChIP-qPCR analyses show a significant increase in the recruitment of Pcf11 and Rhn1 beyond the CPS of *psu1^+^*gene in *ubp8Δ* cells. **f**, ChIP-qPCR analyses demonstrate a significant reduction of Pol II CTD phosphorylation at Tyr1 and Ser2 in *htb1-K119R* cells. **g**, ChIP-qPCR analyses illustrate little to no changes in Tyr1 and Ser2 phosphorylation of Pol II CTD in *ubp8Δ* cells. For panels **c**-**g**, experiments were performed with three independent biological replicates (*n* = 3); one-sided error bars represent the standard deviation (s.d.) of mean qRT-PCR values. *p*-values (Student’s t-test) are indicated in the plots. **h**, A model for elongation to termination suggests that proper levels of H2Bub1 cause Cdk9 to dissociate from the elongation complex. This dissociation allows the recruitment of cleavage and polyadenylation factor (CPF)-associated Dis2, which dephosphorylates Spt5, signaling Pol II to slow down. Slow Pol II leads to the accumulation of Pol II CTD phosphorylation (pTyr1 and pSer2), which facilitates the recruitment of termination factor, Pcf11 and Rhn1. However, when H2Bub1 is absent, Cdk9 remains associated with the elongation complex, preventing the recruitment of CPF-Dis2. As a result, high levels of phosphorylated Spt5 (pSpt5) persist, promoting active elongation and leading to read-through transcription. The CPS is marked by dotted lines.

Interestingly, Cdk9 inhibition in *htb1-K119R* cells narrowed the Pol II distribution beyond the CPS, indicating a rescue of the termination defects observed in these cells (Fig. 4a,b, Extended Data Fig 7). This result suggests that the accumulation of Cdk9 following the loss of H2Bub1 is the primary trigger for the termination defects by disrupting Dis2 recruitment and subsequently stabilizing Spt5 phosphorylation. To investigate the termination defects further, we performed regular PCR and RT-qPCR for selected regions upstream and downstream of the CPS. The DNA gel and RT-qPCR showed transcript production beyond the canonical termination zone of *psu1^+^*, *pgk1^+^*, *pyk1^+^* and *pho1+* genes in *htb1-K119R* cells, implying readthrough transcription (Fig. c, Extended Data Fig 8).

In investigating the molecular mechanisms of termination defects upon disruption of H2Bub1, we conducted ChIP for the termination factors, Pcf11, and Rhn1 (Rtt103). ChIP-qPCR showed a significant reduction in the occupancy of the termination factors Rhn1 and Pcf11 past the CPS in *htb1-K119R* cells (Fig. 4d, Extended Data Fig. 9a,c). However, in Ubp8-deficient backgrounds, there was a marked increase in the chromatic occupancy of Rhn1 and Pcf11 beyond the CPS (Fig. 4e, Extended Data Fig. 9b,d). Thus, the impairment in the recruitment of termination factors can explain the termination defects *in htb1-K119R* cells. During termination, the phosphorylation of Tyr1 (pTyr1) and Ser2 (pSer2) within Pol II CTD heptad repeats assists in recruiting mRNA 3’-end processing and termination factors ^21–24^. The ChIP-qPCR results indicate a substantial decrease in both pTyr1 and pSer2 levels following the disruption of H2Bub1 in *htb1-K119R* cells (Fig. 4f, Extended Data Fig 10a,b,e,f), with only minimal or no changes observed in *ubp8Δ* cells (Fig. 4g, Extended Data Fig. 10c,d,g,h). Consistently, immunoblot revealed a significant decrease in pTyr1 and pSer2 levels in whole cell extract upon the loss of H2Bub1; however, these marks remained unchanged following the stabilization of H2Bub1 (Extended Data Fig. 10i). Consequently, the changes observed in the chromatin occupancy of Rhn1 and Pcf11 in both *htb1-K119R* and *ubp8Δ* strains seem to result directly from the altered levels of pTyr1 and pSer2 on chromatin. In conclusion, our findings suggest that hyper- monoubiquitylation ensures faithful termination by: (i) facilitating the dissociation of Cdk9 from the elongation complex as Pol II traverses the CPS, thus (ii) increasing Dis2’s occupancy and the subsequent dephosphorylation of Spt5, (iii) enhancing the recruitment of cleavage and polyadenylation factors for proper 3’-end processing of mRNA, (iv) subsequently increasing the phosphorylation levels of Tyr1 and Ser2 on the Pol II CTD, which in turn facilitates (v) the chromatin recruitment of termination factors (Fig. 4h; top panel). Actually, in physiological conditions, H2Bub1 accumulates just upstream of the CPS, where pSpt5 gradually decays, explaining its decisive role in the elongation-to-termination transition (Extended Data Fig. 10j) ^16^. However, the loss of H2Bub1 leads to the accumulation of Cdk9 around the CPS, reversing all downstream sequential events and resulting in termination defects (Fig. 4h; bottom panel). Our findings highlight the need for further investigation to uncover the molecular mechanisms by which H2Bub1 differentially regulates Cdk9’s chromatin occupancy throughout the transcription cycle.

The major roles of H2Bub1 in transcription regulation have been depicted in the stages of initiation and elongation. However, the direct role of H2Bub1 in regulating the transition from elongation to termination has not been reported earlier. Therefore, identifying the role of H2Bub1 in termination is novel and adds a new layer of mechanism in transcription regulation. It would be interesting in the future to investigate how H2Bub1 regulates Cdk9 occupancy, pTyr1, and pSer2 during the elongation to termination transition. Does H2Bub1 influence the recruitment of the kinase(s) and/or phosphatase(s) of pTyr1 and pSer2? Several mechanistic explanations for the role of H2Bub1 in transcription have already been proposed. H2Bub1 stimulates FACT-mediated displacement of an H2A/H2B dimer from the core nucleosome, thereby enhancing the passage of Pol II through the nucleosome ^25^. H2Bub1 promotes the reassembly of nucleosomes in the wake of elongating Pol II ^26,27^. Moreover, H2Bub1 regulates transcription by facilitating the recruitment of Set1, a component of the COMPASS complex (MLL complex in humans), and by Dot1, which catalyzes the methylation of histone H3 lysine 4 (H3K4me) and 79 (H3K79me), respectively ^28–30^. These two marks are linked to active transcription. Therefore, in the future, it would be interesting to determine whether any of these mechanisms play intermediate roles in regulating the chromatin occupancy of Cdk9 by H2Bub1.

## Acknowledgements

We thank Robert P. Fisher, Jason. C. Tanny B. Schwer, S. Shuman, and V. Vanoosthuyse for providing yeast strains and antibodies; Robert C. Coleman, Matthew J. Gamble and Ian Willis for manuscript reading and providing critical comments. This work was supported by start-up fund from Montefiore-Einstein Cancer Center (MECC), Albert Einstein College of Medicine.

## Author Contributions

T.B. conducted immunoblot analysis and performed ChIP-qPCR analysis. T.B. conducted ChIP-seq and data analysis to map genome-wide distribution of Cdk9, Pch1, Spt5, pSpt5, Pol II, and Dis2. B.B. isolated RNA from yeast and conducted RT-qPCR analysis. P.K.P and T.B. generated yeast strains. P.K.P. and T.B. prepared the manuscript.

## Author Information

Correspondence and requests for materials should be addressed to

## METHODS

### Yeast strains and growth conditions

Yeast strains used in the study are listed in Extended Data Table 1. Strains were grown in YES (5% yeast extract, 30% dextrose, 250 mg/l each of adenine, leucine, lysine, histidine and uracil) media at 30°C^31,32^, unless otherwise indicated. New strains were generated by mating or random spore analysis^33,34^. Strains with epitope tags and deletions were generated by homologous recombination of DNA fragments either synthesized by PCR or fragmented from a donor plasmid^35,36^. Correct marker integration was confirmed by PCR.

### Cell lines and media

Colon carcinoma-derived HCT116 cells were cultured in McCoy’s 5A medium with L-glutamine (Corning) supplemented with 10% Fetal Bovine Serum (FBS, Gibco) and 1x Penicillin-Streptomycin (Corning).

### Immunoblotting

Antibodies used in this study recognized Pol II (BioLegend, Clone: CTD4H8, Catalog No: 904004), Pol II CTD phospho Ser2 (Abcam; Catalog No. Ab5095), Pol II CTD phospho Tyr1 (Active Motif; Catalog No. 61383), H2B (Active Motif; Catalog No. 39237), H2Bub1(Active Motif; Catalog No. 39623) and phospho Spt5 or total Spt5 (*S. pombe*) (Dr. Larochelle (21st Century Biochemicals; Pr1357) ^8^. For immunoblot analysis, proteins were separated by SDS-PAGE and transferred to Protran^TM^ 0.45 μm nitrocellulose membranes (Cytiva Amersham™). The membranes were probed with primary antibodies at dilutions recommended by the suppliers. Immunoblots were developed with Horseradish Peroxidase (HRP) conjugated donkey anti-rabbit (GE Healthcare Life Sciences; Catalog No. NA934V), sheep anti-mouse (GE Healthcare Life Sciences; Catalog No. NA9310V) and Alexa Fluor-coupled goat anti-rabbit (A21076, Life Technologies). Proteins were detected by enhanced chemiluminescence (Pierce ECL Western Blotting Substrate, Thermo Scientific; Catalog No. 32106).

### Chromatin immunoprecipitation in fission yeast

Chromatin immunoprecipitation (ChIP) was performed as previously described ^7,16^.. Briefly, *S. pombe* cell cultures were grown in YES to an O.D_600_ of 0.5 – 0.6 and crosslinked with 1% formaldehyde for 15 minutes at room temperature. Crosslinking was terminated by adding 2.5 M glycine to a final concentration of 125 mM for 5 minutes. Cells were pelleted by centrifugation, washed with cold 1x TBS (10 mM Tris pH 7.5, 150 mM NaCl), frozen on dry ice, and stored at -80°C. Cell pellets from 50 ml cultures were resuspended in ChIP lysis buffer I (50 mM HEPES pH 7.5, 140 mM NaCl, 1 mM EDTA, 1% Triton X-100, 0.1% Na-deoxycholate, 1 mM PMSF, and protease and phosphatase inhibitor cocktail) and lysed in a Mini-Beadbeater-16 (BioSpec Products; Catalog No. 607) in the presence of glass beads (0.1 mm diameter) (BioSpec Products; Cat. No. 11079101) at 4°C for 1 minute (with 1 minute intervals on ice). Lysates were centrifuged at 16,100 x g for 20 minutes at 4°C. Pellets containing chromatin were resuspended in 1 ml ChIP lysis buffer II (50 mM HEPES pH 7.5, 140 mM NaCl, 1 mM EDTA, 1% Triton X-100, 0.1% SDS, 0.1% Na-deoxycholate, 1 mM PMSF, and protease and phosphatase inhibitor cocktail) and transferred to 15 ml polycarbonate tubes. Lysates were sonicated for 30 minutes at 4°C (30 seconds ON, 30 seconds OFF, output setting high) using a water bath sonicator (Diagenode Bioruptor® Plus; Cat. No. B01020001), transferred to new 1.5 ml tubes, and centrifuged at 16,100 x g for 20 minutes at 4°C. 250 – 300 µg of chromatin extract was incubated with antibodies, FLAG (SIGMA; Catalog No. F3165- 1MG), Myc (Clone: 9E10; BioLegend; Catalog No. 626802), Pol II (8WG16) (Biolegend; Catalog No. 664906), Pol II CTD phospho Ser2 (Abcam; Catalog No. Ab5095), Pol II CTD phospho Tyr1 (Active Motif; Catalog No. 61383), V5 (PK) (Fortis Life Sciences; Catalog No. A190-120A), H2B (Active Motif; Catalog No. 39237), H2Bub1(Active Motif; Catalog No. 39623), H3 (Active Motif; Catalog No. 39763), phospho Spt5 ^8^ (*S. pombe*) (Dr. Larochelle (21st Century Biochemicals; Pr1357), overnight at 4°C with constant nutation. Extracts from *S. cerevisiae* strain BY4742 were also prepared in parallel and spiked into each immunoprecipitation (1% of the weight) for normalization. The suspension was incubated at room temperature for an additional 1.5 – 2 hours with Dynabeads Protein G beads (Invitrogen™; Cat. No. 10004D), preblocked with 1 mg/ml BSA and 0.3 mg/ml Salmon Sperm DNA (Invitrogen™; Catalog No. AM9680). The beads were then washed 2x with ChIP lysis buffer I, 2x with high salt ChIP wash buffer (50 mM HEPES pH 7.5, 500 mM NaCl, 1 mM EDTA, 1% Triton X-100, 0.1% Na-deoxycholate), 2x with ChIP wash buffer (10 mM Tris-HCl pH 8.0, 0.25 M LiCl, 0.5% NP-40, 0.5% Na-deoxycholate, 1 mM EDTA), and 1x with 1x TE buffer (10 mM Tris-HCl pH 8.0, 1 mM EDTA). After washing, the suspension was centrifuged at 1700 x g for 20 – 30 seconds. Protein-DNA complexes were eluted from the beads with 1x ChIP elution buffer (50 mM Tris-HCl pH 8.0, 1% SDS, 1mM EDTA) by incubating at 65°C for 20 – 30 minutes with occasional vortexing. To reverse the protein-DNA crosslinks, the suspension was incubated overnight at 65°C with 0.2 M NaCl. The non-crosslinked suspension was treated with 1 µg of RNase A (Thermo Scientific™; Catalog No. FEREN0531) at 37°C for 30 minutes and with 0.8 units of Proteinase K (Thermo Scientific™; Catalog No. FEREO0491) at 45°C for 45 minutes. The DNA was purified using the QIAquick PCR Purification Kit (Qiagen; Catalog No. 28106) according to the manufacturer’s protocol. The purified DNA was subjected to either qPCR with Radiant™ Green qPCR Master Mix (2x) (Alkali Scientific; Catalog No. QS1001) in 384-well plates or library preparation for sequencing.

### Chromatin immunoprecipitation in human cells

ChIP-qPCR experiments were done as described previously ^37^. Briefly, HCT116 cells grown to 50-60% confluence were crosslinked with 1% formaldehyde for 10 minutes at 25°C. Crosslinking was quenched with 125 mM glycine for 5 minutes at 25°C. Cells were washed twice with ice-cold PBS and collected into 1 ml of RIPA (150 mM NaCl, 1% v/v Nonidet P-40, 0.5% w/v deoxycholate, 0.1% w/v SDS, 50 mM Tris pH 8.0, 5 mM EDTA) buffer supplemented with protease and phosphatase inhibitors for each 150- mm dish. Cells were lysed, and chromatin was sheared by sonication in a Bioruptor at high power, for 3 x 10 minutes with cycles of 30 seconds ON and 30 seconds OFF. Lysates were clarified by centrifugation at 20,000 x g for 20 minutes at 4°C. Before immunoprecipitation, lysates (∼5 x 10^6 cells per experiment) were pre-cleared with Pierce™ Protein A Agarose (Thermo Scientific™; Catalog No. PI20333) for 2 hours at 4°C. Beads were separated by centrifugation at 4000 x g for 1 minute at 4°C. The resulting supernatant was incubated with anti- Cdk9 antibody (Abclonal; Catalog No. A11145) and Pol II (D8L4Y) (Cell Signaling Technology; Catalog No. 14958S) for 4 hours at 4°C with constant nutation. The suspension was incubated at 25°C for an additional 2 hours with Dynabeads Protein G beads, pre-blocked with 1 mg/ml BSA and 0.25 mg/ml Salmon Sperm DNA. The beads were washed 2x with RIPA buffer, 4x with Szak IP wash buffer (100 mM Tris-HCl, pH 8.5, 500 mM lithium chloride, 1% (v/v) nonidet-P-40, 1% (w/v) sodium deoxycholate), 2x with RIPA buffer, and 2x with TE buffer. After all wash steps, centrifugation was performed at 1700 x g for 1 minute at 4°C. Protein-nucleic acid complexes were eluted from the beads with elution buffer (46 mM Tris-HCl, pH 8.0, 0.65 mM EDTA, 1% SDS) by incubating at 65°C for 15 minutes with occasional vortexing. Crosslinking was reversed by incubating at 65°C for 16 hours. The un-crosslinked suspension was treated with 1 μg of RNase A at 37°C for 30 minutes and with 0.8 units of Proteinase K at 45°C for 45 minutes. DNA was purified using the QIAquick® PCR Purification Kit according to the manufacturer’s protocol. The purified DNA was subjected to either qPCR or library preparation for sequencing.

### ChIP-seq analysis

Multiplexed ChIP-seq libraries were prepared using the NEBNext® Ultra™ II DNA Library Preparation kit (E7103S) with 25-50 ng of input or IP DNA and barcode adaptors (NEBNext Multiplex Oligos for Illumina [Set 1, E7335] & [Set 2, E7500]). 150-nucelotides (150- nt) paired-end sequencing (PE150) was performed on NovaSeq 6000. Quality control and adapter trimming of FASTQ (raw sequencing) files were done in Galaxy using FastQC Read Quality reports (Galaxy Version 0.74) and Trimmomatic flexible read trimming tool for Illumina NGS data (Galaxy Version 0.39), respectively. Processed FASTQ files were aligned to the *S. pombe* or human genome (version hg19) using Bowtie2^38^ (Galaxy Version 2.5.3). Aligned sequences of each biological replicate were fed into MACS2^39^ (Galaxy Version 2.2.9.1) to call peaks from alignment results. Generated “bedgraph treatment” files were converted to bigwig using “Convert BedGraph to BigWig” (Galaxy Version 1.0.1), mean signal was computed using “bigwigCompare” (Galaxy Version 3.5.4). Generated bigwig files were processed using computeMatrix (Galaxy Version 3.5.4) in DeepTools^40^ to prepare data for plotting heatmaps and/or profiles of given regions. Heatmap and Metagene plots were generated using “plot Heatmap” (Galaxy Version 3.5.4) and “plotProfile” (Galaxy Version 3.5.4) tools, respectively. The log_2_ signal ratios were calculated using “bigwigCompare” (Galaxy Version 3.5.4).

### Quantitative RT-PCR (RT-qPCR)

*S. pombe* cell cultures (50 mL) of strains JS78 (WT), LV259 (*htb1-K119R*), and TB4 (*ubp8D*) were grown to OD_600_ of 0.5. Total RNA was extracted by a hot phenol method ^41^ and further purified on Qiagen RNeasy columns. For strand-specific RT-qPCR to measure levels of *psu1^+^*, *pgk1^+^*, *pyk1^+^* and *pho1^+^* transcripts, 1 μg total RNA was converted to cDNA using the superscript III kit (Invitrogen) and gene-specific primers listed in Extended Data Table 2. Expression levels were normalized to those of the 18sDNA transcript in each strain.

## Data Access

The raw and processed sequencing files have been submitted to the NCBI Gene Expression Omnibus (GEO; https://www.ncbi.nlm.nih.gov/geo/query/acc.cgi) (Accession number: GSE271342). The published data used in this study under the following accession code: GSE115682 ^16^ and GSE138548 ^42^.

---

**Extended Data Figure 1.**
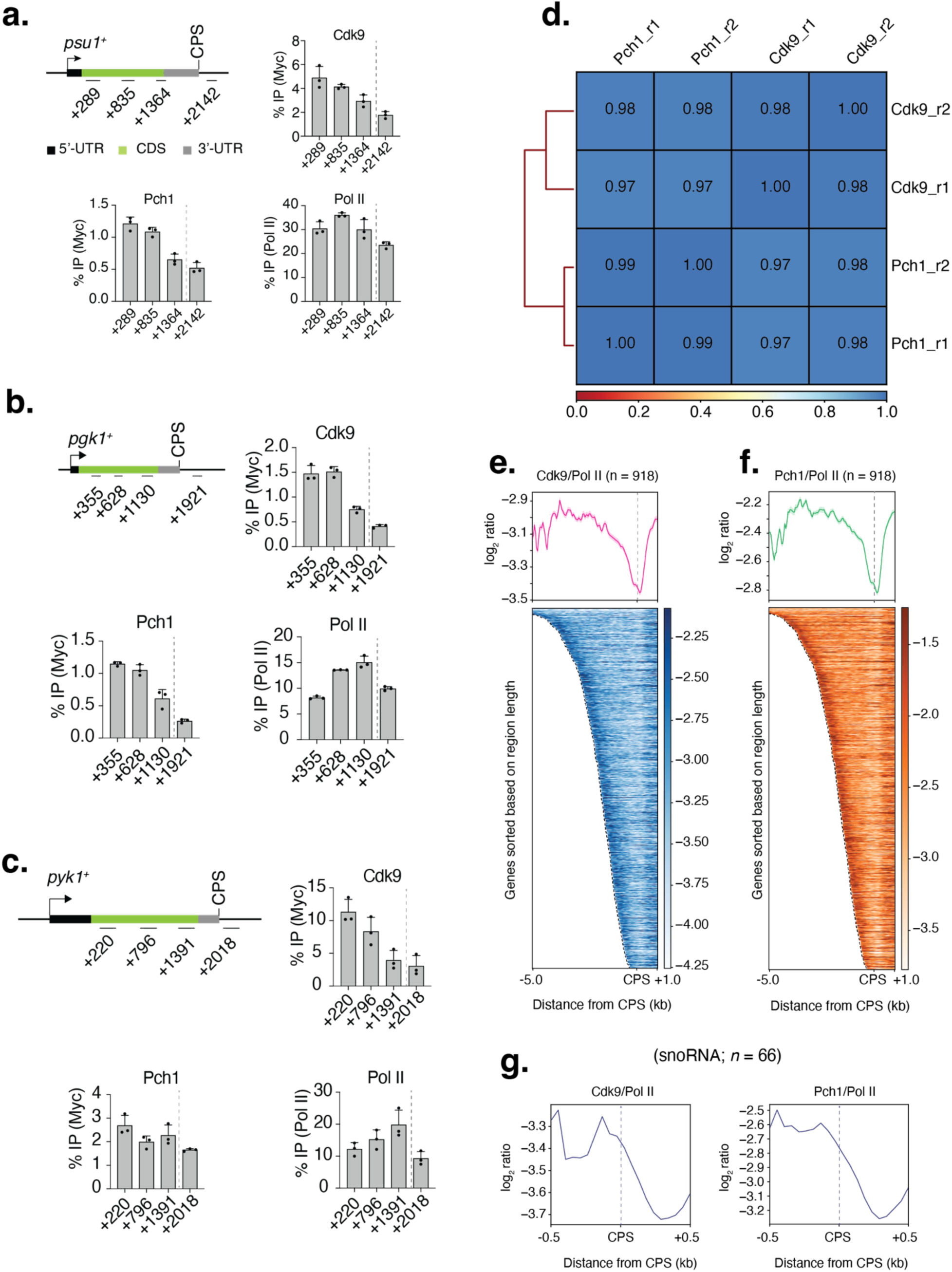
Release of Cdk9 and Pch1 from the elongation complex in fission yeast when Pol II traverses the CPS. **a**, ChIP-qPCR analyses show a gradual decrease in the occupancy of Cdk9 and Pch1 towards the 3’-end of the *psu1^+^*gene. **b**, ChIP-qPCR analyses show a gradual decrease in the occupancy of Cdk9 and Pch1 towards the 3’- end of the *pgk1^+^* gene. **c**, ChIP-qPCR analyses show a gradual decrease in the occupancy of Cdk9 and Pch1 towards the 3’-end of the *pyk1^+^* gene. For panels **a**-**c**, individual data values were obtained from three (*n* = 3) independent experiments; one-sided error bars represent the standard deviation (s.d.) of mean ChIP values. **d**, Correlation between ChIP-seq samples. Paired-end sequencing reads were mapped to the fission yeast genome using Bowtie2 (Galaxy Version 2.5.3+galaxy1). Mapped reads of each biological replicate were used to calculate the correlation between pairs of replicates. Values in boxes represent the Pearson correlation coefficient between corresponding samples (*n* = 2 biological replicates). **e**, Genome-wide distribution of Cdk9. Metagene analyses (top) and heatmaps (bottom) show a comparison between log_2_ ratios of Cdk9:Pol II for Pol II-transcribed genes (1 kb separated; *n* = 918). **f**, Genome-wide distribution of Pch1. Metagene analyses (top) and heatmaps (bottom) show a comparison between log_2_ ratios of Pch1:Pol II for Pol II-transcribed genes (1 kb separated; *n* = 918). For panels **e** and **f**, genes were sorted based on region length between -5 kb to +1 kb relative to the CPS. **g**, Distribution of Cdk9 and Pch1 on snoRNA genes. Metagene analyses show genome-wide comparison between log_2_ ratios of Cdk9:Pol II or Pch1:Pol II for Pol II-transcribed snoRNA genes (*n* = 66) from -0.5 kb to +0.5 kb relative to the CPS. The CPS is marked by vertical dotted lines.

**Extended Data Figure 2.**
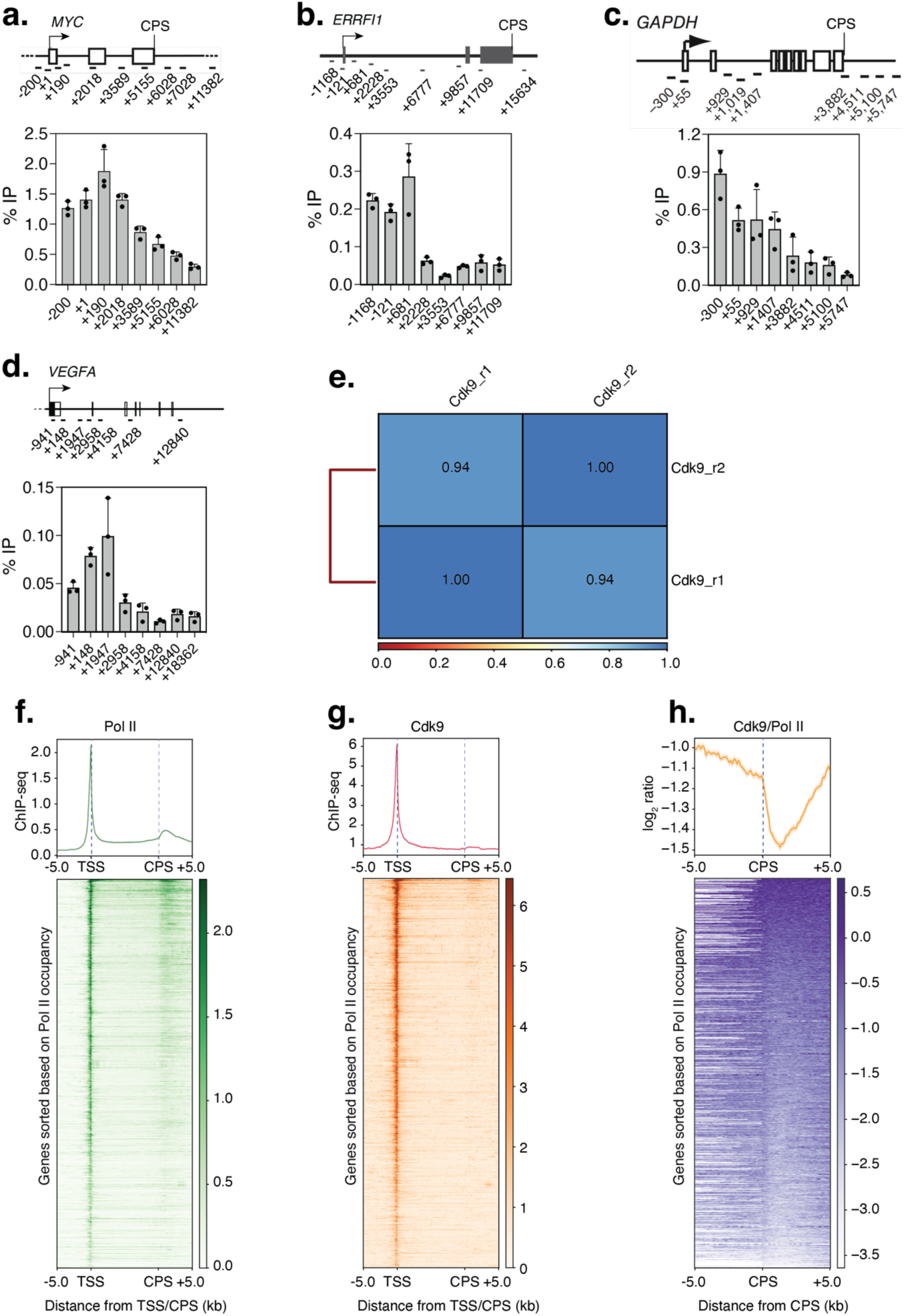
Cdk9 dissociation as Pol II crosses the CPS in human cells. **a**-**d**, ChIP-qPCR analyses show a gradual decrease in Cdk9 occupancy towards the 3’-end of *MYC* (**a**), *ERRFI1* (**b**), *GAPDH* (**c**), and *VEGFA* (**d**). Individual data values in panels **a**-**d** were obtained from three independent experiments (*n* = 3). One-sided error bars represent the standard deviation (s.d.) of the mean ChIP values. **e**, Correlation between Cdk9 ChIP-seq samples. Paired-end sequencing reads were mapped to the human genome (hg19) using Bowtie2 (Galaxy Version 2.5.3+galaxy1). Mapped reads of each biological replicate were used to calculate the correlation between pairs of replicates. Values in boxes represent the Pearson correlation coefficient between corresponding samples (*n* = 2 biological replicates). **f**, Genome-wide distribution of Pol II. Metagene analyses (top) and heatmaps (bottom) show the distribution of Pol II. Pol II ChIP data utilized in this analysis were obtained from a published study ^42^. **g**, Genome-wide distribution of Cdk9. Metagene analyses (top) and heatmaps (bottom) show the distribution of Cdk9. **h**, Genome-wide distribution of Cdk9 with a CPS-centered focus. Metagene analyses (top) and heatmaps (bottom) display the distribution of Cdk9 on Pol II-transcribed genes from -5.0 kb to +5.0 kb. The log2 ratios of Cdk9:Pol II were plotted. For **f**, **g**, the regions between the TSS and the CPS have been scaled for comparison among genes of different lengths. For **f**-**h**, plots were generated using 10 kb separated Pol II-transcribed genes (*n* = 7723). The TSS and CPS are marked by vertical dotted lines.

**Extended Data Figure 3.**
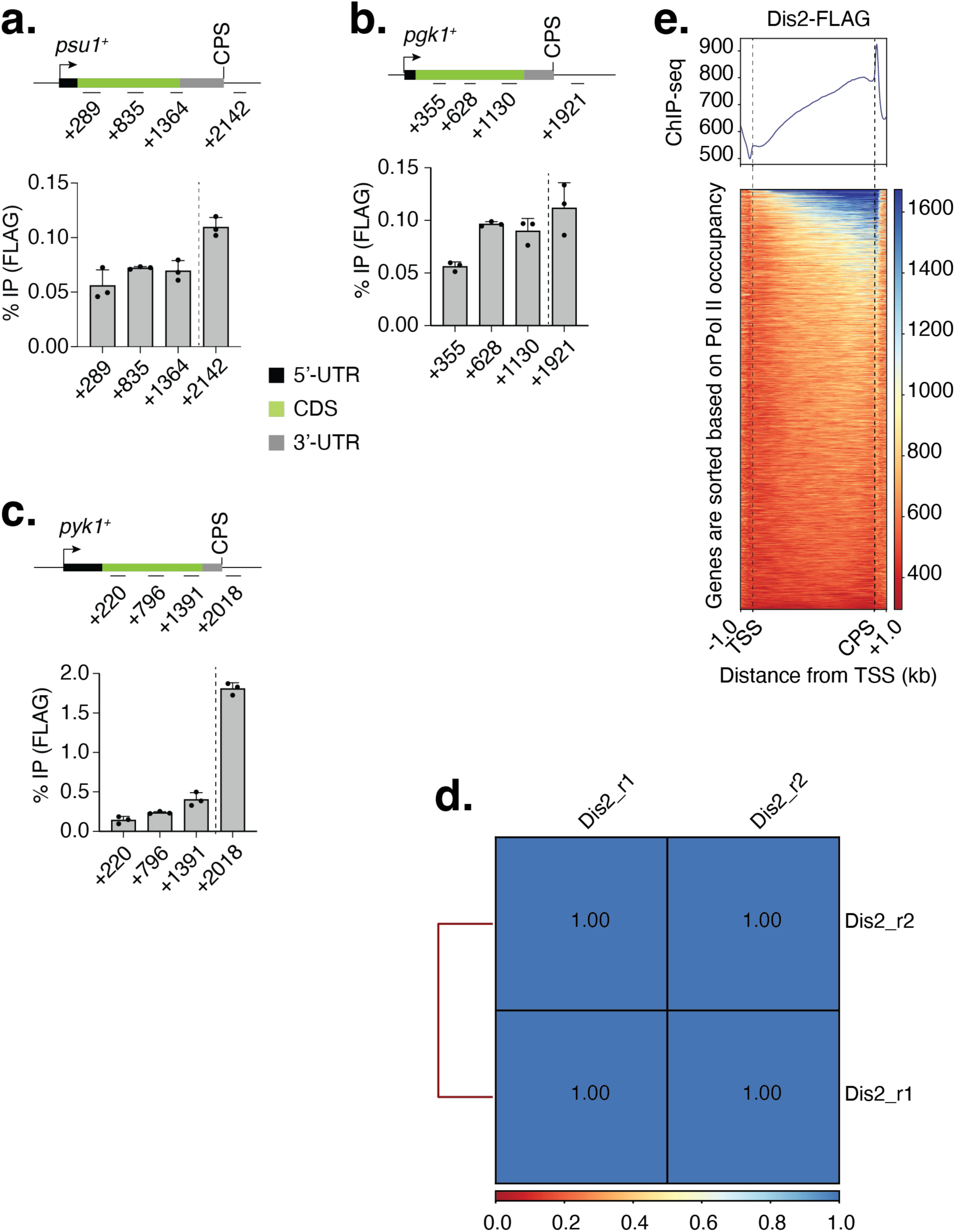
Sharp accumulation of Dis2 after the CPS in fission yeast genes. **a**-**c**, ChIP-qPCR analyses show a gradual increase in Dis2 occupancy towards the 3’-end of *psu1^+^* (**a**), *pgk1^+^* (**b**), and *pyk1^+^* (**c**) genes, with a sharp increase in occupancy after the CPS (marked by vertical dotted lines). Experiments were performed with three biological replicates (*n* = 3). Individual data points are shown in the plots. One-sided error bars represent the standard deviation (s.d.) of the mean ChIP values. **d**, Correlation between Dis2-FLAG ChIP-seq samples. Paired-end sequencing reads were mapped to the fission yeast genome using Bowtie2 (Galaxy Version 2.5.3+galaxy1). Mapped reads of each biological replicate were used to calculate the correlation between pairs of replicates. Values in boxes represent the Pearson correlation coefficient between corresponding samples (*n* = 2 biological replicates). **e**, Genome-wide distribution of Dis2. Metagene analyses (top) and heatmaps (bottom) demonstrate the dramatic accumulation of Dis2 beyond the CPS of Pol II-transcribed genes (*n* = 7723). Genes are sorted based on Pol II occupancy. The regions between the TSS and the CPS have been scaled for comparison among genes of different lengths. The TSS and CPS are marked by vertical dotted lines.

**Extended Data Figure 4.**
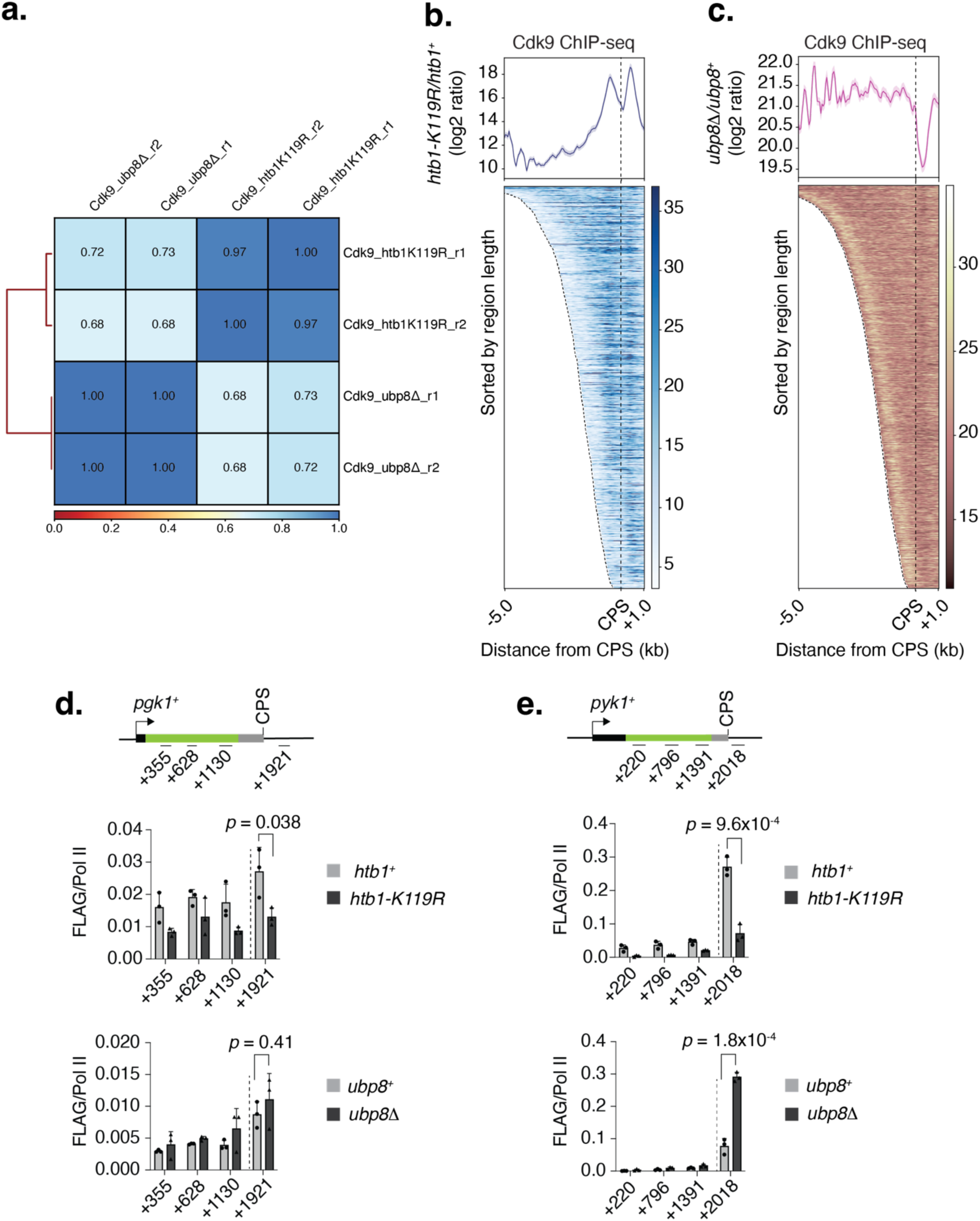
H2Bub1 regulates the distribution of Cdk9 and Dis2. **a**, Correlation between ChIP-seq samples. Paired-end sequencing reads were mapped to the fission yeast genome using Bowtie2 (Galaxy Version 2.5.3+galaxy1). The correlation between pairs of replicates was calculated using mapped reads from each biological replicate. Values in boxes represent the Pearson correlation coefficient between corresponding samples (*n* = 2 biological replicates). **b**, Genome-wide distribution of Cdk9 upon loss of H2Bub1. Metagene analyses (top) and heatmaps (bottom) compare the log_2_ ratios of mean ChIP values in *htb1-K119R* and *htb1^+^* for Pol II-transcribed genes (*n* = 918). **c**, Genome-wide distribution of Cdk9 under persistent levels of H2Bub1. Metagene analyses (top) and heatmaps (bottom) compare the log_2_ ratios of mean ChIP values in *ubp8D* and *ubp8^+^*for Pol II-transcribed genes (*n* = 3329). For **b**, **c**, genes were sorted based on region length between -5 kb to +1 kb relative to the CPS. **d**, Distribution of Dis2 on *pgk1^+^* gene. ChIP-qPCR analyses show a significant decrease in Dis2 occupancy after the CPS (marked by a vertical dotted line) in *htb1-K119R* cells (top panel), with a modest increase in *ubp8D* cells (bottom panel). **e**, Distribution of Dis2 on *pyk1^+^* gene. ChIP-qPCR analyses reveal a significant decrease in Dis2 occupancy after the CPS (marked by a vertical dotted line) in *htb1-K119R* cells (top panel). Conversely, Dis2 occupancy is increased in *ubp8D* cells (bottom panel). For **d**,**e**, experiments were performed with three independent biological replicates (*n* = 3); one-sided error bars show the standard deviation (s.d.) of mean ChIP values; *p*-values (Student’s t-test) are indicated in the plots; Y-axis show ratio plots of Dis2 ChIP over Pol II.

**Extended Data Figure 5.**
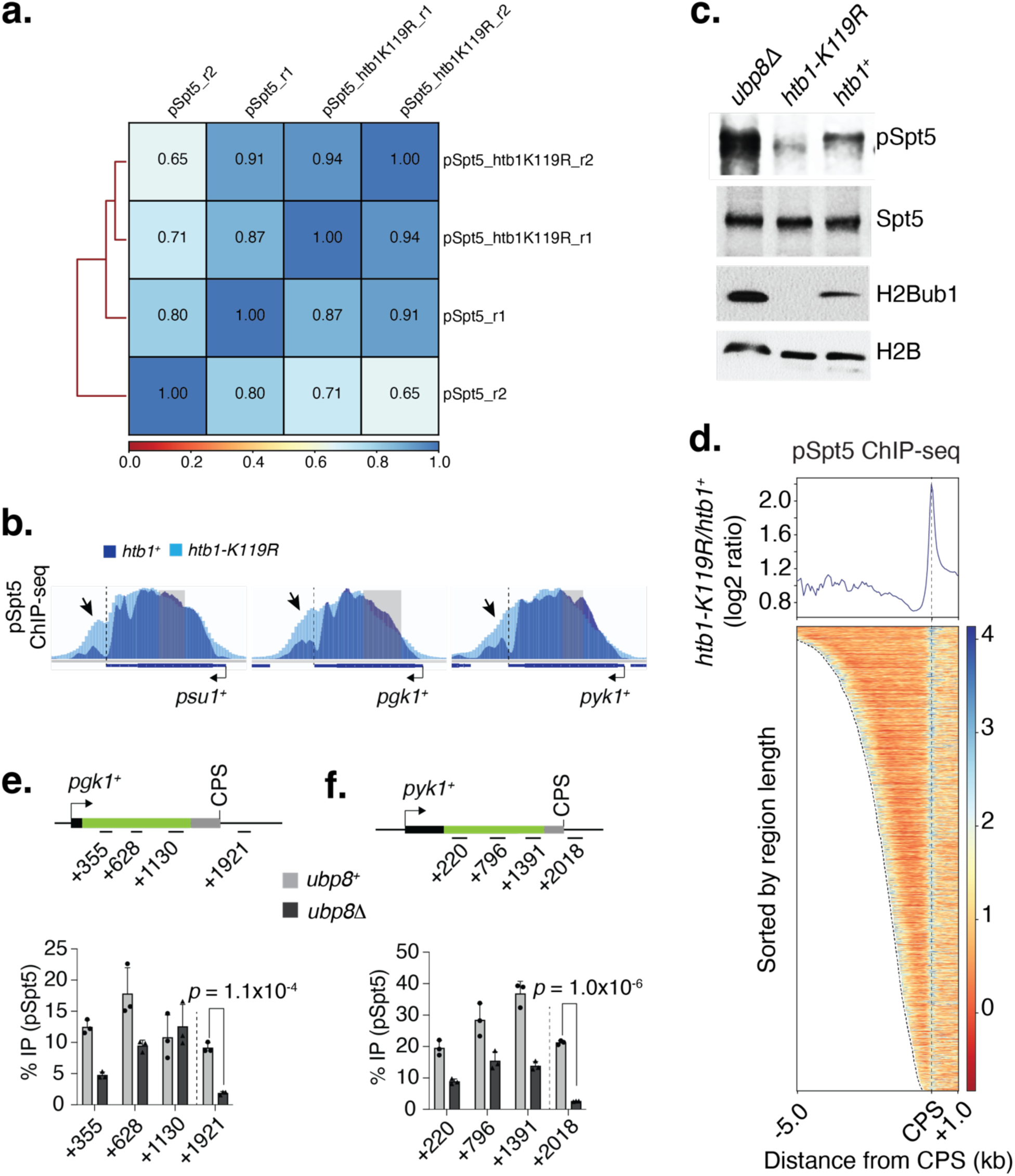
H2Bub1 regulates Spt5 phosphorylation on chromatin. **a**, Correlation between ChIP-seq samples. Paired-end sequencing reads were mapped to the fission yeast genome using Bowtie2 (Galaxy Version 2.5.3+galaxy1). The correlation between pairs of replicates was calculated using mapped reads from each biological replicate. Values in boxes represent the Pearson correlation coefficient between corresponding samples (*n* = 2 biological replicates). **b**, Individual browser tracks display an increase in pSpt5 occupancy downstream of the CPS (marked by an arrow) and a decrease at the 5’ end (marked by a gray shaded area) of *psu1^+^*, *pgk1^+^* and *pyk1^+^*upon loss of H2Bub1. Blue and cyan tracks represent the pSpt5 distribution in *htb1^+^* and *htb1-K119R* cells, respectively, are overlaid. **c**, The immunoblot shows a reduction in pSpt5 in the whole-cell extract of *htb1-K119R*, whereas it is slightly stabilized in *ubp8Δ* cells. Additionally, there is a loss of H2Bub1 in *htb1-K119R* and a slight increase in H2Bub1 in *ubp8Δ* cells. **d**, Genome-wide distribution of pspt5 upon loss of H2Bub1. Metagene analyses (top) and heatmaps (bottom) compare the log_2_ ratios of mean pSpt5 ChIP values in *htb1-K119R* and *htb1^+^* for Pol II-transcribed genes (*n* = 918). The genes were sorted based on region length between -5 kb to +1 kb relative to the CPS. **e**, ChIP-qPCR analyses show the distribution of pSpt5 on *pgk1^+^* gene in *ubp8^+^*(gray bar) and *ubp8Δ* (black bar) cells. Results indicate a significant decrease in pSpt5 occupancy after the CPS in *ubp8Δ* cells. **f**, ChIP-qPCR analyses show the distribution of pSpt5 on *pyk1^+^* gene in *ubp8^+^* (gray bar) and *ubp8Δ* (black bar) cells. Results indicate a significant decrease in pSpt5 occupancy after the CPS in *ubp8Δ* cells. For **e**,**f**, experiments were performed with three independent biological replicates (*n* = 3); one-sided error bars show the standard deviation (s.d.) of mean ChIP values; *p*-values (Student’s t-test) are indicated in the plots. The vertical dotted lines mark the CPS.

**Extended Data Figure 6.**
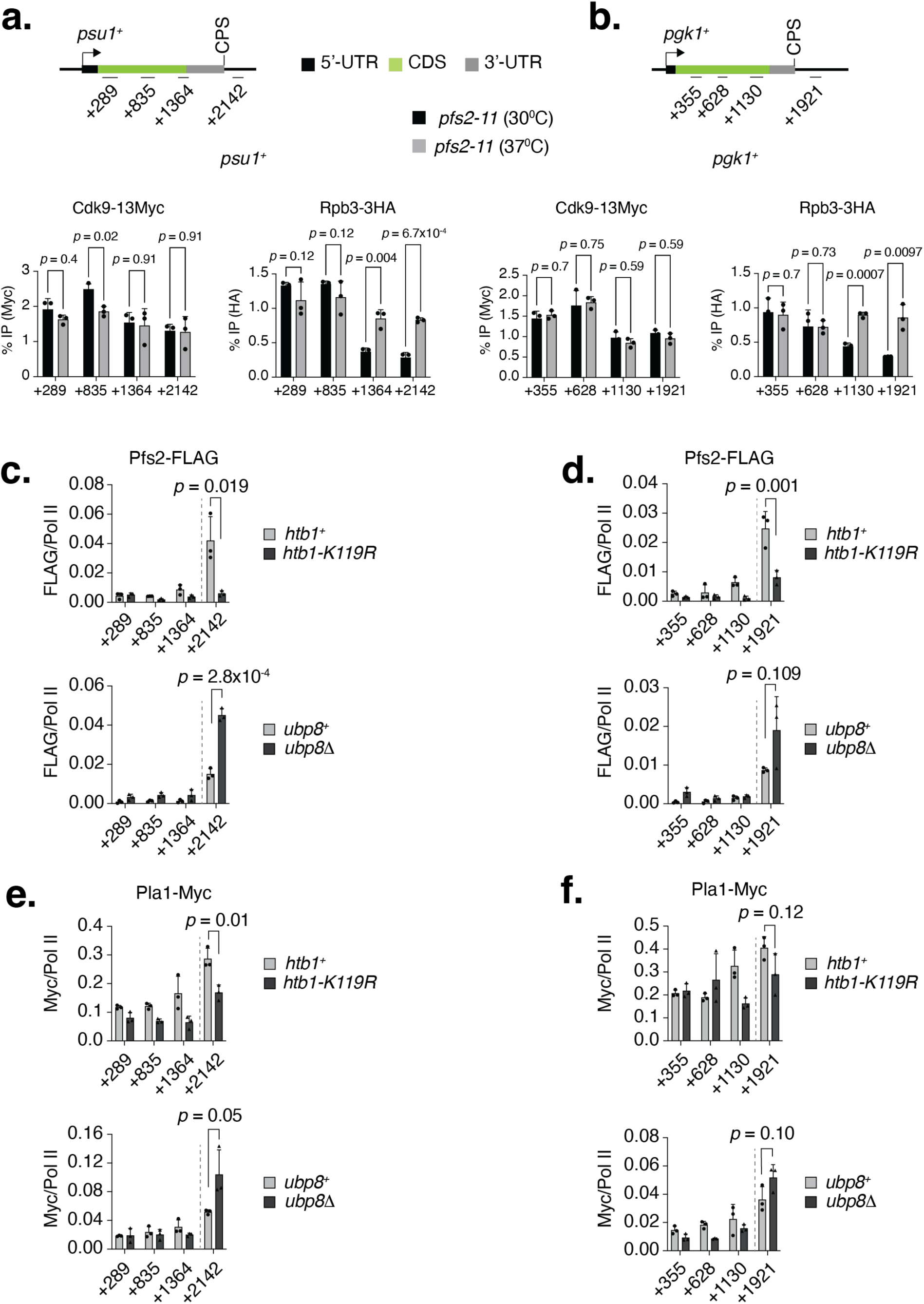
Cdk9 evictions occur upstream of cleavage and polyadenylation. **a**, Chromatin distribution of Cdk9, where ChIP-qPCR results reveal no alterations in Cdk9 occupancy on *psu1^+^* upon heat inactivation of Pfs2 at 37°C, while Pol II occupancy on *psu1^+^* increased at the 3’-end of the gene under the same condition. **b**, ChIP-qPCR analyses demonstrate no changes in Cdk9 occupancy on *pgk1^+^*upon heat inactivation of Pfs2 at 37°C, with a concurrent increase in Pol II occupancy at the 3’-end of the gene of *pgk1^+^*. **c**, ChIP-qPCR results indicating a decrease in Pfs2 occupancy on *psu1^+^* in *htb1-K119R* cells (top panel) and an increase in Pfs2 occupancy in *ubp8Δ* cells (bottom panel). **d**, ChIP-qPCR results show a parallel pattern for *pgk1^+^*, with decreased Pfs2 occupancy in *htb1-K119R* cells (top panel) and increased occupancy in *ubp8Δ* cells (bottom panel). **e**, ChIP-qPCR results demonstrating a decrease in Pla1 occupancy on *psu1^+^* in *htb1-K119R* cells (top panel) and an increase in *ubp8Δ* cells (bottom panel). **f**, ChIP-qPCR reveals a decrease in Pla1 occupancy on *pgk1^+^* in *htb1-K119R* cells (top panel) and an increase in *ubp8Δ* cells (bottom panel). For panels **a**-**f**, individual data points were obtained from three independent biological replicates (*n* = 3). One-sided error bars represent the standard deviation (s.d.) of mean ChIP values, and *p*-values (Student’s t-test) are indicated in the plots. Vertical dotted lines mark the CPS.

**Extended Data Figure 7.**
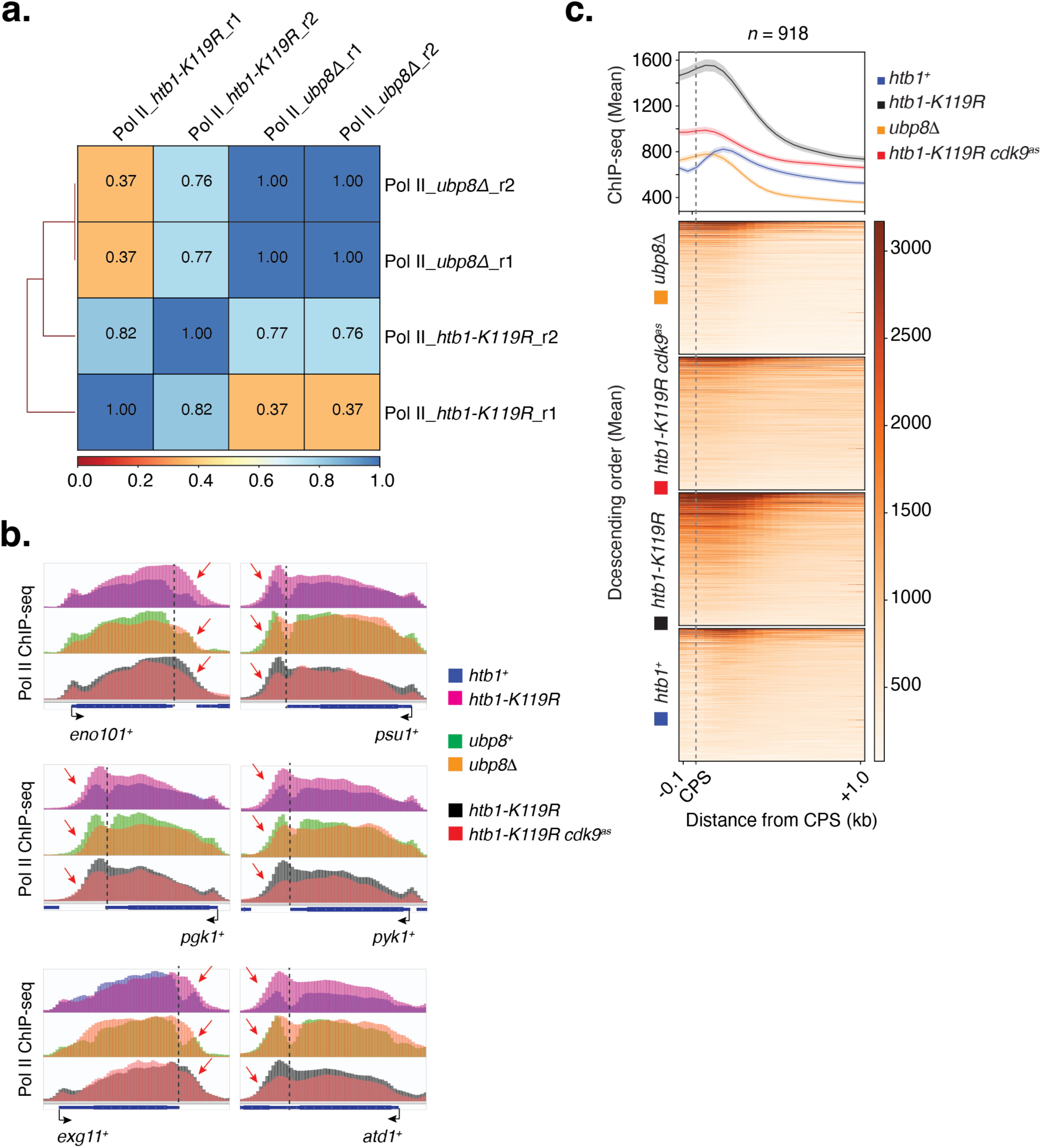
H2Bub1 regulates transcription termination. **a**, Correlation between Pol II ChIP-seq samples in *htb1-K119R* and *ubp8Δ* cells. Paired-end sequencing reads were mapped to the fission yeast genome using Bowtie2 (Galaxy Version 2.5.3+galaxy1). The correlation between pairs of replicates was calculated using mapped reads from each biological replicate. Values in boxes represent the Pearson correlation coefficient between corresponding samples (*n* = 2 biological replicates). **b**, Loss of H2Bub1 impairs termination. ChIP-seq browser tracks reveal an extended Pol II distribution in *htb1-K119R* cells beyond the regular termination zone seen in *htb1^+^* cells, indicating readthrough transcription, i.e., termination defects. The blue and pink tracks, representing the Pol II distribution in *htb1^+^* and *htb1-K119R* cells, respectively, are overlaid. Conversely, persistent levels of H2Bub1 favor termination. ChIP-seq browser tracks show a narrowed Pol II distribution in the termination zone in *ubp8Δ* cells, indicating that hyper mono-ubiquitylation favors termination. The green and orange tracks, representing the Pol II distribution in *ubp8^+^* and *ubp8Δ* cells, respectively, are overlaid. Cdk9 inhibition in the *htb1-K119R* background narrows Pol II distribution, suggesting a rescue of termination defects. This indicates that the accumulation of Cdk9 following the loss of H2Bub1 is the main trigger for termination defects by continuing elongation. The black and red tracks, representing the Pol II distribution in *htb1-K119R*, with or without inhibition of Cdk9, respectively, are overlaid. Pol II ChIP-seq data following Cdk9 inhibition in *htb1-K119R* cells, used in this analysis, were obtained from a published study ^16^. The regions of alterations in Pol II distribution after the CPS are indicated by red arrows. **c**, Metagene analyses (top) and heatmaps (bottom) display a genome-wide distribution of Pol II beyond the CPS of 1 kb separated genes (*n* = 918) in *htb1-K119R* cells, both with and without Cdk9 inhibition, as well as in *ubp8Δ* cells. Metagene plots further support global termination defects in *htb1-K119R* cells, proper termination upon hyper-monoubiquitylation in *ubp8Δ* cells, and the rescue of termination defects in *htb1-K119R* cells upon Cdk9 inhibition. The genes are sorted based on Pol II occupancy between regions -0.1 kb to +1.0 kb of the CPS. For **b**,**c**, the vertical dotted lines mark the CPS. The Cdk9 inhibition ChIP-seq data was used from previous report ^16^.

**Extended Data Figure 8.**
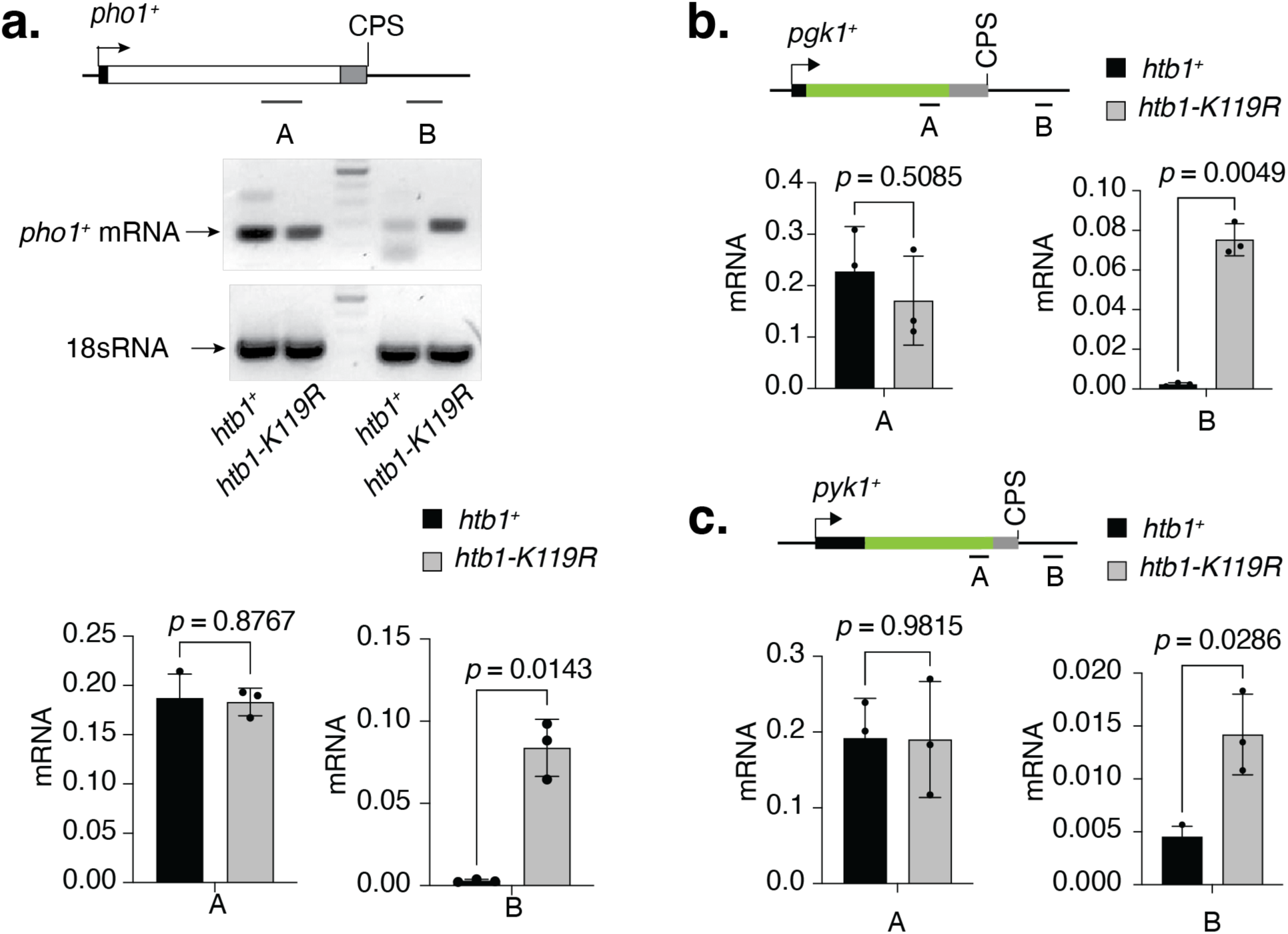
H2Bub1 regulates transcription termination. **a**, The top panel shows an agarose gel visualizing transcript production from the upstream (A) and downstream (B) regions of the CPS of the *pho1^+^*gene. The results indicate the synthesis of *pho1^+^* mRNA beyond the regular termination zone in *htb1-K119R* cells, with little to no mRNA synthesis in *htb1^+^*cells. The bottom panel shows RT-qPCR analyses quantifying *pho1^+^*mRNA production from the same regions, A and B, indicating increased mRNA synthesis beyond the termination zone in *htb1-K119R* cells. **b**, RT-qPCR analyses quantifying *pgk1^+^* mRNA production from the regions, A and B, indicating increased mRNA synthesis beyond the termination zone in *htb1-K119R* cells. **c**, RT-qPCR analyses quantifying *pyk1^+^* mRNA production from the regions, A and B, indicating increased mRNA synthesis beyond the termination zone in *htb1-K119R* cells. For panels **a**-**c**, individual data points shown in the plots were obtained from three independent biological replicates (*n* = 3); two-sided error bars represent the standard deviation (s.d.) of mean qRT-PCR values; *p*-values (Student’s t-test) are indicated in the plots.

**Extended Data Figure 9.**
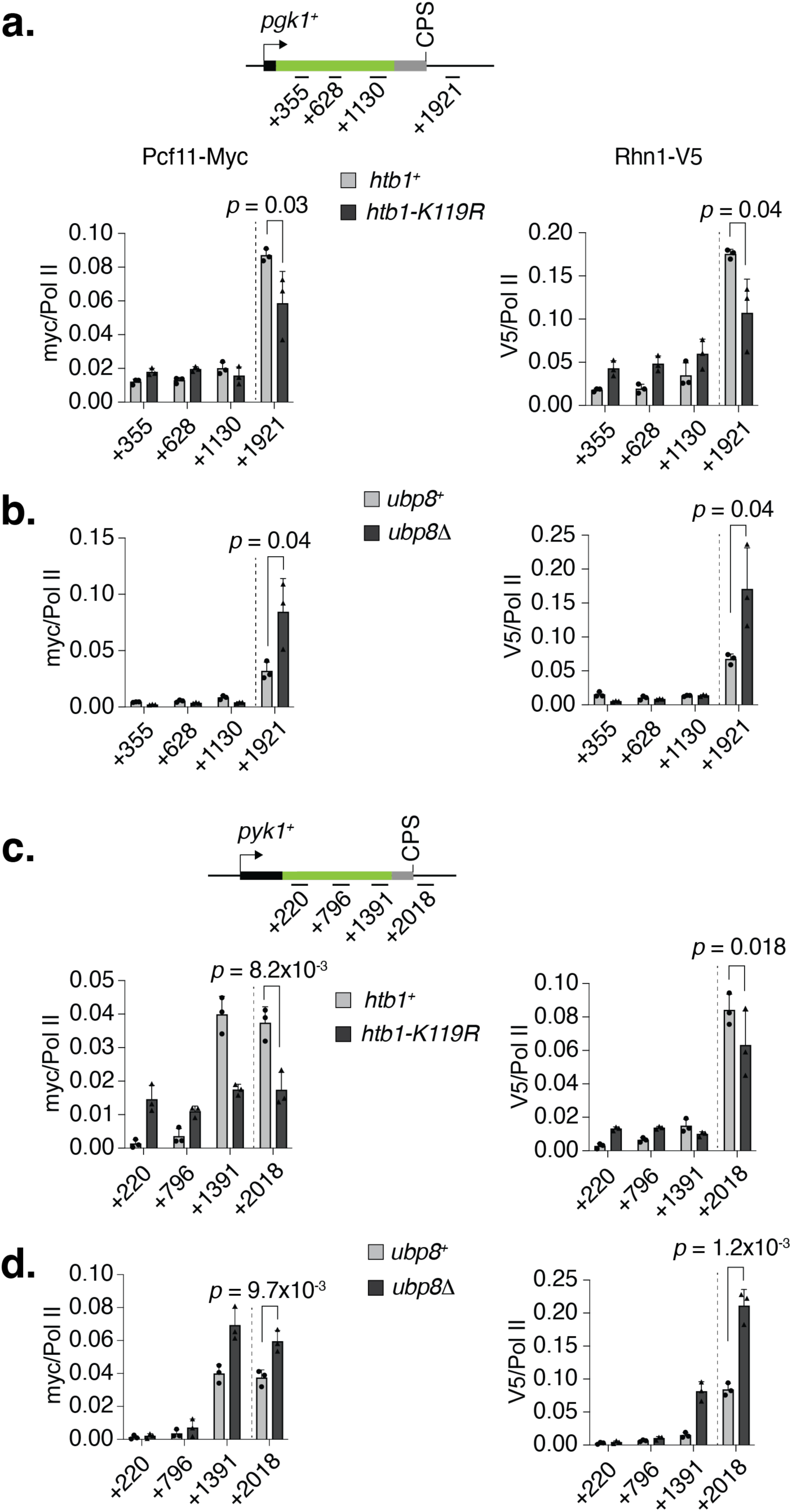
H2Bub1 controls termination factor recruitment. **a**, Distribution of Pcf11 and Rhn1 on *pgk1^+^*. ChIP-qPCR analyses show a significant decrease in the occupancy of Pcf11 and Rhn1 in *htb1-K119R* cells. **b**, ChIP-qPCR analyses show a significant increase in the occupancy of Pcf11 and Rhn1 in *ubp8Δ* cells. **c**, Distribution of Pcf11 and Rhn1 on *pyk1^+^*. ChIP-qPCR analyses show a significant decrease in the occupancy of Pcf11 and Rhn1 in *htb1-K119R* cells. **d**, ChIP-qPCR analyses show a significant increase in the occupancy of Pcf11 and Rhn1 in *ubp8Δ* cells. For panels **a**-**d**, experiments were performed with three independent biological replicates (*n* = 3); one-sided error bars represent the standard deviation (s.d.) of mean ChIP values. *p*-values (Student’s t-test) are indicated in the plots. The CPS is marked by dotted lines.

**Extended Data Figure 10.**
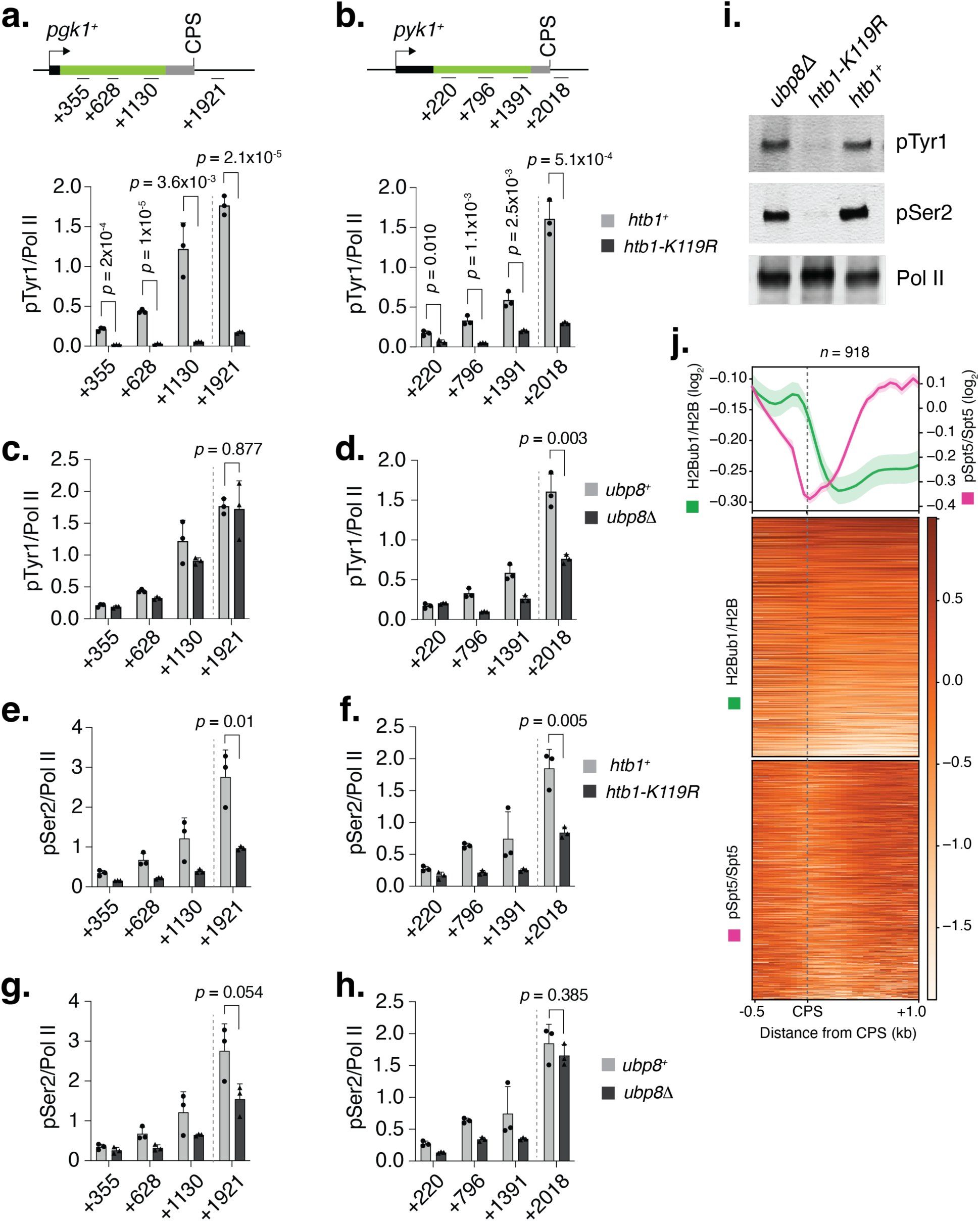
H2Bub1 influences Pol II CTD Tyr1 and Ser2 phosphorylation. Distribution of Pol II CTD Tyr1 and Ser2 phosphorylation on *pgk1^+^* and *pyk1^+^*. **a**, **b**, ChIP-qPCR analyses indicate a significant reduction in Pol II CTD Tyr1 phosphorylation in *htb1-K119R* cells. **c**, **d**, Panels reveal minimal to no change in Pol II CTD Tyr1 phosphorylation in *ubp8Δ* cells, as demonstrated by ChIP-qPCR analyses. **e**, **f**, Similarly, illustrate a notable decrease in Pol II CTD Ser2 phosphorylation in *htb1-K119R* cells through ChIP-qPCR analyses. **g**, **h**, Panels confirm that *ubp8Δ* cells exhibit little to no change in Pol II CTD Ser2 phosphorylation according to ChIP-qPCR analyses. All experiments (panels **a**-**h**) were conducted with three independent biological replicates (*n* = 3); one-sided error bars represent the standard deviation (s.d.) of mean ChIP values, and *p*-values (Student’s t-test) are shown in the plots. The CPS is marked by dotted lines. **i**, Immunoblot showing a loss of Pol II CTD Tyr1 (pTyr1) and Ser2 (pSer2) phosphorylation in whole cell extracts of *htb1-K119R* cells, whereas these modifications remain nearly unchanged in *ubp8Δ* cells. **j**, Metagene analyses (top) and heatmaps (bottom) display a genome-wide distribution of pSpt5:Spt5 and H2Bub1:H2B around the CPS of 1 kb separated genes (*n* = 918). The results indicate the accumulation of H2Bub1 upstream of the CPS, where phospho-Spt5 gradually decays. The genes are sorted based on Pol II occupancy between regions -0.5 kb to +1.0 kb of the CPS. H2B, H2Bub1, Spt5, pSpt5 ChIP-seq data used in this analysis were obtained from a published study ^16^.

**Extended Data Table 1.** Yeast Strains.

| Strain | Genotype | Source |
| --- | --- | --- |
| JS78 | *leu1-32 ura4-D18 his3-D1 ade6-M210 h* | <sup>43</sup> |
| LV239 | *htb1-K119R-FLAG::kanMX6 spt5-13myc::kanMX6 leu1-32 ura4-D18 his3-D1 ade6-M21X h?* | <sup>8</sup> |
| LV259 | *htb1-K119R-FLAG::kanMX6 cdk9-13myc::kanMX6 leu1-32 ura4-D18 his3-D1 ade6-M210 h?* | <sup>8</sup> |
| HD6-51 | *cdk9-13myc::kanMX6 leu1-32 ura4-D18 his3-D1 ade6-M210 h+* | <sup>44</sup> |
| CS112 | *cdk9T<sup>120G</sup>::kanMX6 spt5-13myc::kanMX6 leu1-32 ura4-D18 his3-D1 ade6-* | <sup>41</sup> |
| CS172 | *pch1-13myc::kanMX6 leu1-32 ura4-D18 his3-D1 ade6-M216 h* | <sup>45</sup> |
| VV1484 | *dis2-3xFLAG::natMX6 h-* | <sup>46</sup> |
| TB4 | *h+ cdk9-13myc::kanMX6 ubp8Δ::ura4* | This article |
| TB11 | *rhn1-PK::hph pcf11-myc::kanMX6 htb1-K119R::kanMX6 h?* | This article |
| TB21 | *dis2-3xFLAG::natMX6 htb1-K119R::kanMX6* | This article |
| TB27 | *dis2-3xFLAG::natMX6 ubp8Δ::ura4* | This article |
| TB29 | *rhn1-PK::hph pcf11-myc::kanMX6 ubp8Δ::ura4 h?* | This article |
| TB32 | *pla1-myc::kanMX6 ubp8Δ::ura4* | This article |
| TB38 | *pfs2-11 cdk9-myc::kanMX6 h+* | This article |
| TB54 | *cdk9as::natMX6 htb1K119R::kanMX6 pfs2-FLAG::kanMX6* | This article |
| TB55 | *pla1-myc::kanMX6 htb1K119R::kanMX6* | This article |
| TB57 | *pfs2-FLAG::kanMX6 ubp8Δ::ura4* | This article |
| FY23510 | *h90 leu1-32 ura4-D18 ade6-M210 red1-FLAG::kanMX6 pla1-myc::kanMX6* | Yeast Genetic Resource Center (YGRC), Japan |
| FY23525 | *h90 leu1-32 ura4-D18 ade6-M210 or M216 rhn1-PK::hph pcf11-myc::kanMX6* | Yeast Genetic Resource Center (YGRC), Japan |
| FY23541 | *h90 or h- leu1-32 ura4-D18 ade6-210 red5-myc::hph pfs2-FLAG::kanMX6* | Yeast Genetic Resource Center (YGRC), Japan |

**Extended Data Table 2.**
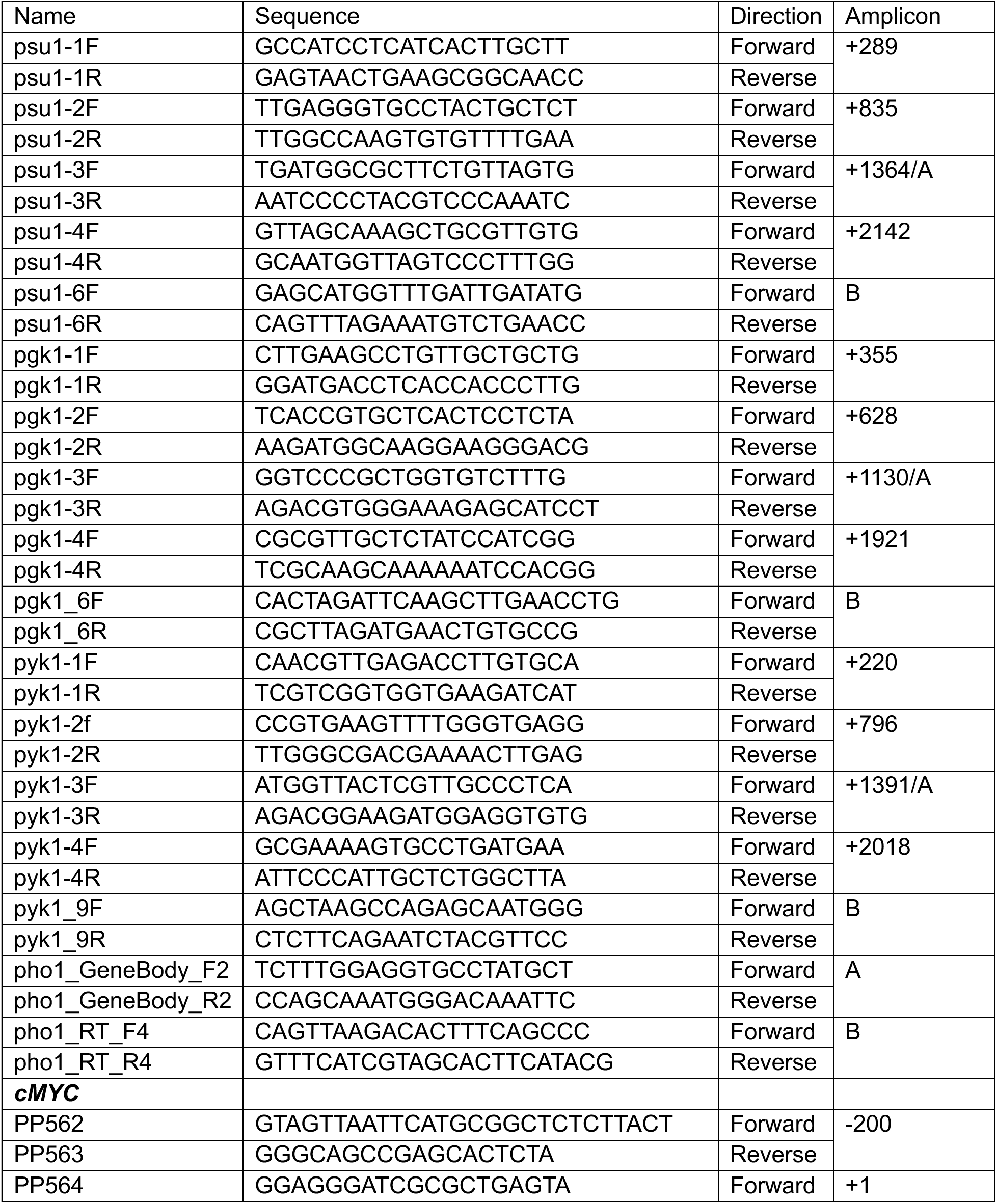
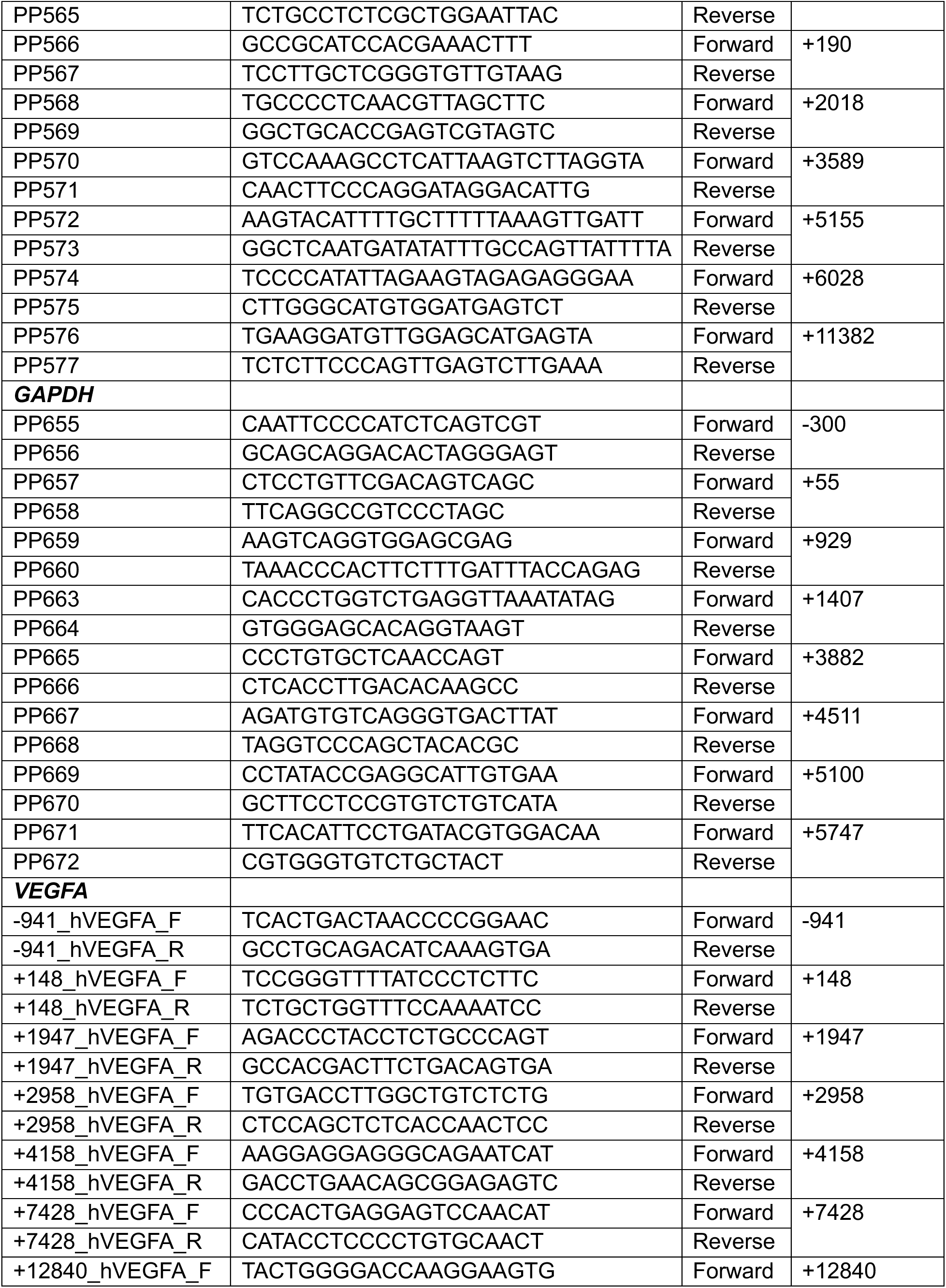
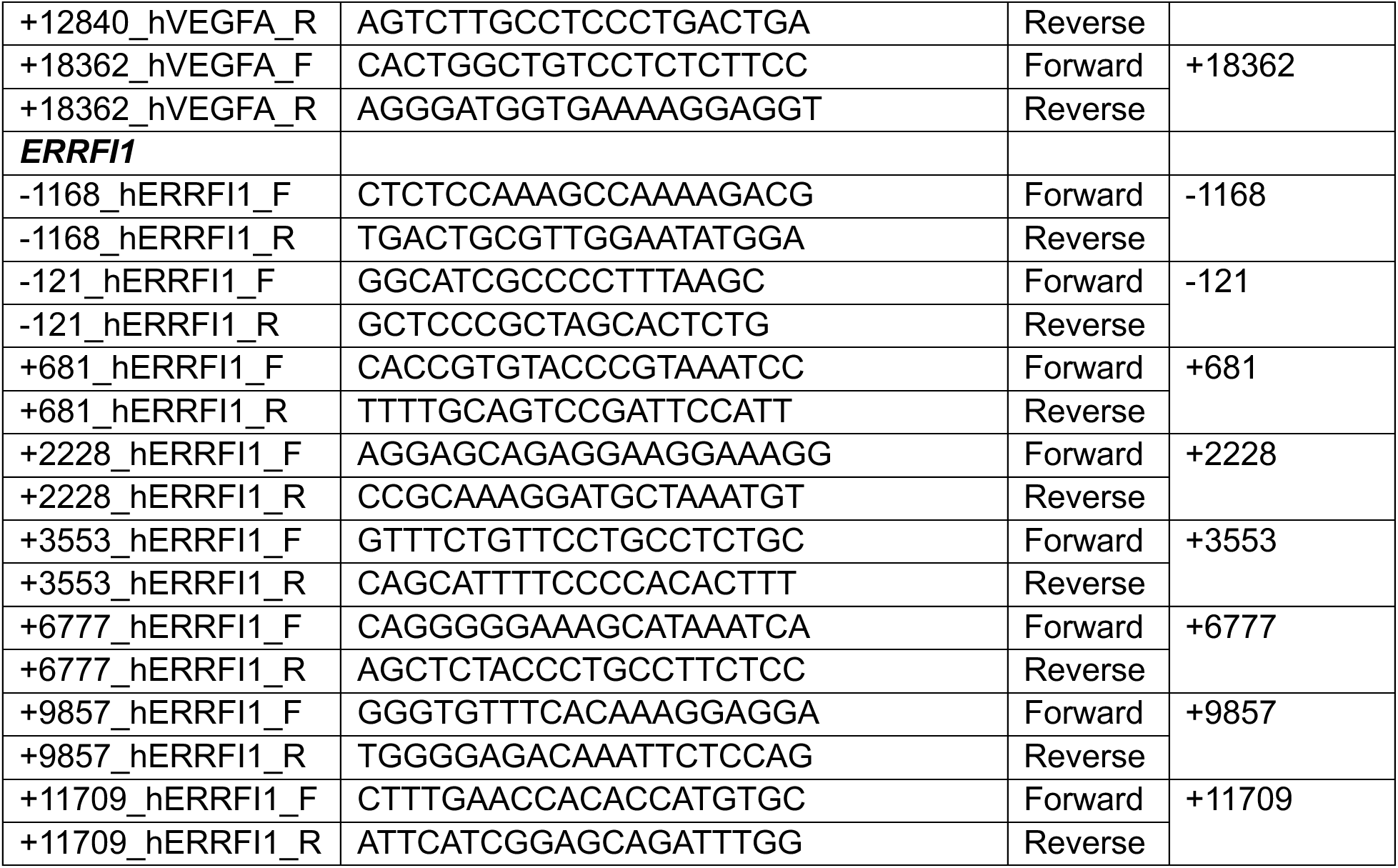
Oligonucleotides.

## References

1 Parua, P. K. & Fisher, R. P. Dissecting the Pol II transcription cycle and derailing cancer with CDK inhibitors. Nat Chem Biol 16, 716–724, doi:10.1038/s41589-020-0563-4 (2020).

2 Proudfoot, N. J. Transcriptional termination in mammals: Stopping the RNA polymerase II juggernaut. Science 352, aad9926, doi:10.1126/science.aad9926 (2016).

3 Rodriguez-Molina, J. B., West, S. & Passmore, L. A. Knowing when to stop: Transcription termination on protein-coding genes by eukaryotic RNAPII. Mol Cell 83, 404–415, doi:10.1016/j.molcel.2022.12.021 (2023).

4 Grallert, A. et al. A PP1-PP2A phosphatase relay controls mitotic progression. Nature 517, 94–98, doi:10.1038/nature14019 (2015).

5 Zofall, M. & Grewal, S. I. HULC, a histone H2B ubiquitinating complex, modulates heterochromatin independent of histone methylation in fission yeast. J Biol Chem 282, 14065–14072, doi:10.1074/jbc.M700292200 (2007).

6 Helmlinger, D. et al. The S. pombe SAGA complex controls the switch from proliferation to sexual differentiation through the opposing roles of its subunits Gcn5 and Spt8. Genes Dev 22, 3184–3195, doi:10.1101/gad.1719908 (2008).

7 Parua, P. K. et al. A Cdk9-PP1 switch regulates the elongation-termination transition of RNA polymerase II. Nature 558, 460–464, doi:10.1038/s41586-018-0214-z (2018).

8 Sanso, M. et al. A positive feedback loop links opposing functions of P-TEFb/Cdk9 and histone H2B ubiquitylation to regulate transcript elongation in fission yeast. PLoS Genet 8, e1002822, doi:10.1371/journal.pgen.1002822 (2012).

9 Wood, A., Schneider, J., Dover, J., Johnston, M. & Shilatifard, A. The Bur1/Bur2 complex is required for histone H2B monoubiquitination by Rad6/Bre1 and histone methylation by COMPASS. Mol Cell 20, 589–599, doi:10.1016/j.molcel.2005.09.010 (2005).

10 Laribee, R. N. et al. BUR kinase selectively regulates H3 K4 trimethylation and H2B ubiquitylation through recruitment of the PAF elongation complex. Curr Biol 15, 1487–1493, doi:10.1016/j.cub.2005.07.028 (2005).

11 Zhou, K., Kuo, W. H., Fillingham, J. & Greenblatt, J. F. Control of transcriptional elongation and cotranscriptional histone modification by the yeast BUR kinase substrate Spt5. Proc Natl Acad Sci U S A 106, 6956–6961, doi:10.1073/pnas.0806302106 (2009).

12 Liu, Y. et al. Phosphorylation of the transcription elongation factor Spt5 by yeast Bur1 kinase stimulates recruitment of the PAF complex. Mol Cell Biol 29, 4852–4863, doi:10.1128/MCB.00609-09 (2009).

13 Qiu, H., Hu, C., Gaur, N. A. & Hinnebusch, A. G. Pol II CTD kinases Bur1 and Kin28 promote Spt5 CTR-independent recruitment of Paf1 complex. EMBO J 31, 3494–3505, doi:10.1038/emboj.2012.188 (2012).

14 Mayekar, M. K., Gardner, R. G. & Arndt, K. M. The recruitment of the Saccharomyces cerevisiae Paf1 complex to active genes requires a domain of Rtf1 that directly interacts with the Spt4-Spt5 complex. Mol Cell Biol 33, 3259–3273, doi:10.1128/MCB.00270-13 (2013).

15 Minsky, N. et al. Monoubiquitinated H2B is associated with the transcribed region of highly expressed genes in human cells. Nat Cell Biol 10, 483–488, doi:10.1038/ncb1712 (2008).

16 Sanso, M. et al. Cdk9 and H2Bub1 signal to Clr6-CII/Rpd3S to suppress aberrant antisense transcription. Nucleic Acids Res 48, 7154–7168, doi:10.1093/nar/gkaa474 (2020).

17 Bres, V., Gomes, N., Pickle, L. & Jones, K. A. A human splicing factor, SKIP, associates with P-TEFb and enhances transcription elongation by HIV-1 Tat. Genes Dev 19, 1211–1226, doi:10.1101/gad.1291705 (2005).

18 Bres, V., Yoshida, T., Pickle, L. & Jones, K. A. SKIP interacts with c-Myc and Menin to promote HIV-1 Tat transactivation. Mol Cell 36, 75–87, doi:10.1016/j.molcel.2009.08.015 (2009).

19 Boreikaite, V. & Passmore, L. A. 3’-End Processing of Eukaryotic mRNA: Machinery, Regulation, and Impact on Gene Expression. Annu Rev Biochem 92, 199–225, doi:10.1146/annurev-biochem-052521-012445 (2023).

20 Wang, S. W., Asakawa, K., Win, T. Z., Toda, T. & Norbury, C. J. Inactivation of the pre- mRNA cleavage and polyadenylation factor Pfs2 in fission yeast causes lethal cell cycle defects. Mol Cell Biol 25, 2288–2296, doi:10.1128/MCB.25.6.2288-2296.2005 (2005).

21 Descostes, N. et al. Tyrosine phosphorylation of RNA polymerase II CTD is associated with antisense promoter transcription and active enhancers in mammalian cells. Elife 3, e02105, doi:10.7554/eLife.02105 (2014).

22 Larochelle, M. et al. Common mechanism of transcription termination at coding and noncoding RNA genes in fission yeast. Nat Commun 9, 4364, doi:10.1038/s41467-018-06546-x (2018).

23 Ahn, S. H., Kim, M. & Buratowski, S. Phosphorylation of serine 2 within the RNA polymerase II C-terminal domain couples transcription and 3’ end processing. Mol Cell 13, 67–76 (2004).

24 Bartkowiak, B. et al. CDK12 is a transcription elongation-associated CTD kinase, the metazoan ortholog of yeast Ctk1. Genes Dev 24, 2303–2316, doi:10.1101/gad.1968210 (2010).

25 Pavri, R. et al. Histone H2B monoubiquitination functions cooperatively with FACT to regulate elongation by RNA polymerase II. Cell 125, 703–717, doi:10.1016/j.cell.2006.04.029 (2006).

26 Fleming, A. B., Kao, C. F., Hillyer, C., Pikaart, M. & Osley, M. A. H2B ubiquitylation plays a role in nucleosome dynamics during transcription elongation. Mol Cell 31, 57–66, doi:10.1016/j.molcel.2008.04.025 (2008).

27 Batta, K., Zhang, Z., Yen, K., Goffman, D. B. & Pugh, B. F. Genome-wide function of H2B ubiquitylation in promoter and genic regions. Genes Dev 25, 2254–2265, doi:10.1101/gad.177238.111 (2011).

28 Ng, H. H., Xu, R. M., Zhang, Y. & Struhl, K. Ubiquitination of histone H2B by Rad6 is required for efficient Dot1-mediated methylation of histone H3 lysine 79. J Biol Chem 277, 34655–34657, doi:10.1074/jbc.C200433200 (2002).

29 Briggs, S. D. et al. Gene silencing: trans-histone regulatory pathway in chromatin. Nature 418, 498, doi:10.1038/nature00970 (2002).

30 Sun, Z. W. & Allis, C. D. Ubiquitination of histone H2B regulates H3 methylation and gene silencing in yeast. Nature 418, 104–108, doi:10.1038/nature00883 (2002).

31 Forsburg, S. L. Growth and manipulation of S. pombe. Curr Protoc Mol Biol **Chapter** 13, Unit 13 16, doi:10.1002/0471142727.mb1316s64 (2003).

32 Forsburg, S. L. S. pombe strain maintenance and media. Curr Protoc Mol Biol **Chapter** 13, Unit 13 15, doi:10.1002/0471142727.mb1315s64 (2003).

33 Sabatinos, S. A. & Forsburg, S. L. Molecular genetics of Schizosaccharomyces pombe. Methods Enzymol 470, 759–795, doi:10.1016/S0076-6879(10)70032-X (2010).

34 Forsburg, S. L. & Rhind, N. Basic methods for fission yeast. Yeast 23, 173–183, doi:10.1002/yea.1347 (2006).

35 Bahler, J. et al. Heterologous modules for efficient and versatile PCR-based gene targeting in Schizosaccharomyces pombe. Yeast 14, 943–951, doi:10.1002/(SICI)1097-0061(199807)14:10<943::AID-YEA292>3.0.CO;2-Y (1998).

36 Forsburg, S. L. Introduction of DNA into S. pombe cells. Curr Protoc Mol Biol **Chapter** 13, Unit 13 17, doi:10.1002/0471142727.mb1317s64 (2003).

37 Larochelle, S. et al. Cyclin-dependent kinase control of the initiation-to-elongation switch of RNA polymerase II. Nat Struct Mol Biol 19, 1108–1115 (2012).

38 Langmead, B. & Salzberg, S. L. Fast gapped-read alignment with Bowtie 2. Nat Methods 9, 357–359, doi:10.1038/nmeth.1923 (2012).

39 Feng, J., Liu, T., Qin, B., Zhang, Y. & Liu, X. S. Identifying ChIP-seq enrichment using MACS. Nat Protoc 7, 1728–1740, doi:10.1038/nprot.2012.101 (2012).

40 Ramirez, F. et al. deepTools2: a next generation web server for deep-sequencing data analysis. Nucleic Acids Res 44, W160–165, doi:10.1093/nar/gkw257 (2016).

41 Viladevall, L. et al. TFIIH and P-TEFb coordinate transcription with capping enzyme recruitment at specific genes in fission yeast. Mol Cell 33, 738–751, doi:10.1016/j.molcel.2009.01.029 (2009).

42 Parua, P. K., Kalan, S., Benjamin, B., Sanso, M. & Fisher, R. P. Distinct Cdk9- phosphatase switches act at the beginning and end of elongation by RNA polymerase II. Nat Commun 11, 4338, doi:10.1038/s41467-020-18173-6 (2020).

43 Saiz, J. E. & Fisher, R. P. A CDK-activating kinase network is required in cell cycle control and transcription in fission yeast. Curr Biol 12, 1100–1105 (2002).

44 Pei, Y. et al. Cyclin-dependent kinase 9 (Cdk9) of fission yeast is activated by the CDK- activating kinase Csk1, overlaps functionally with the TFIIH-associated kinase Mcs6, and associates with the mRNA cap methyltransferase Pcm1 in vivo. Mol Cell Biol 26, 777–788, doi:10.1128/MCB.26.3.777-788.2006 (2006).

45 St Amour, C. V., et al. Separate domains of fission yeast Cdk9 (P-TEFb) are required for capping enzyme recruitment and primed (Ser7-phosphorylated) Rpb1 carboxyl-terminal domain substrate recognition. Mol Cell Biol 32, 2372–2383, doi:10.1128/MCB.06657-11 (2012).

46 Vanoosthuyse, V. et al. CPF-associated phosphatase activity opposes condensin- mediated chromosome condensation. PLoS Genet 10, e1004415, doi:10.1371/journal.pgen.1004415 (2014).

